# State-of-the-art structural variant calling: What went conceptually wrong and how to fix it?

**DOI:** 10.1101/2021.01.12.426317

**Authors:** Markus Schmidt, Arne Kutzner

## Abstract

Structural variant (SV) calling belongs to the standard tools of modern bioinformatics for identifying and describing alterations in genomes. Initially, this work presents several complex genomic rearrangements that reveal conceptual ambiguities inherent to the SV representations of state-of-the-art SV callers. We contextualize these ambiguities theoretically as well as practically and propose a graph-based approach for resolving them. Our graph model unifies both genomic strands by using the concept of skew-symmetry; it supports graph genomes in general and pan genomes in specific. Instances of our model are inferred directly from seeds instead of the commonly used alignments that conflict with various types of SV as reported here. For yeast genomes, we practically compute adjacency matrices of our graph model and demonstrate that they provide highly accurate descriptions of one genome in terms of another. An open-source prototype implementation of our approach is available under the MIT license at https://github.com/ITBE-Lab/MA.

## Introduction

Structural Variant (SV) calling is a popular technique for discovering and describing genomic differences and rearrangements. Examples of the application of SV calling are phylogenetic analyses and the discovery of genetic disorders. Known techniques for SV calling are: Alignment based approaches that rely on short reads (Illumina) or long reads (PacBio, Oxford Nanopore) [1-4]. Here SV callers exploit one or several types of evidence for SV detection such as: 1) Deviations from the expected distance between paired reads (read-pair based calling), 2) the locations of supplementary alignments for chimeric reads (split-read based calling) and 3) regions with particularly high or low coverage on the reference genome regarding alignments. Alternatively, SV can be detected by comparing assemblies as e.g. in [5-8]. Finally, there are meta-callers [9-11], which try to achieve high accuracy and high recall rates by combining the output of other SV callers. Overviews presenting all these types of SV callers together with their characteristics and their respective performances can be found in [12-15].

Our work starts by reporting shortcomings of state-of-the-art read-pair and split-read based SV calling. Here we identify inevitable ambiguities inherent to the description of genomic rearrangements via atomic SV. As a solution, we introduce a description of genomic rearrangements using skew-symmetric graphs. A highlight of our graph model is a folding scheme for adjacency matrices that unifies forward strand and reverse strand. The general suitability of graphs for the representation of genomic rearrangements is already demonstrated in [16, 17]. Further, we inspect negative side effects of alignments on SV calling that result from their path oriented nature. For circumventing these side effects, our skew-symmetric graphs are computed using seeds merely. In this context, the seeding is extended by a recursive reseeding technique that takes the position of the Dynamic Programming [18, 19] usually incorporated with alignment computations (as e.g. in the aligners Minimap2 [20] and MA [21]). A practical evaluation using the two *Saccharomyces paradoxus* (wild yeast) genomes UFRJ50816 and YPS138 (data provided by [22]) supports the viability of our approach. In this context, we express the genome UFRJ50816 in terms of the genome YPS138 via a graph that is computed either from the assemblies of both genomes or from simulated PacBio reads and Illumina reads. A discussion of our approach emphasizes its suitability with graph genomes and, in this way, with pan genomes.

## Results

### Ambiguities inherent to the description of genomic rearrangements via atomic SV

In the following, we assume that genomic rearrangements are represented by combinations of the following four atomic SV: Deletion, Insertion, Inversion and Duplication. State-of-the-art SV calling attempts to represent genomic rearrangements by nesting these atomic SV. However, this representation scheme comprises inherent ambiguities in two ways: 1) A set of atomic SV can describe multiple reorganizations and, on the contrary, 2) a single genomic rearrangement can be described by multiple sets of atomic SV. Fig. 1 visualizes these ambiguities via examples. In Fig. 1 A), the combination of a duplication and an inversion describes three different genomic outcomes. In Fig. 1 B), a reference *ABCD* is reorganized to 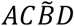 either with two inversions or with a duplication followed by two deletions and an inversion. Recently published works are aware of these ambiguities [23-26]. As an intermediate solution, they exclude complex genomic rearrangements that pose unmanageable ambiguities to them.

**Fig 1.**
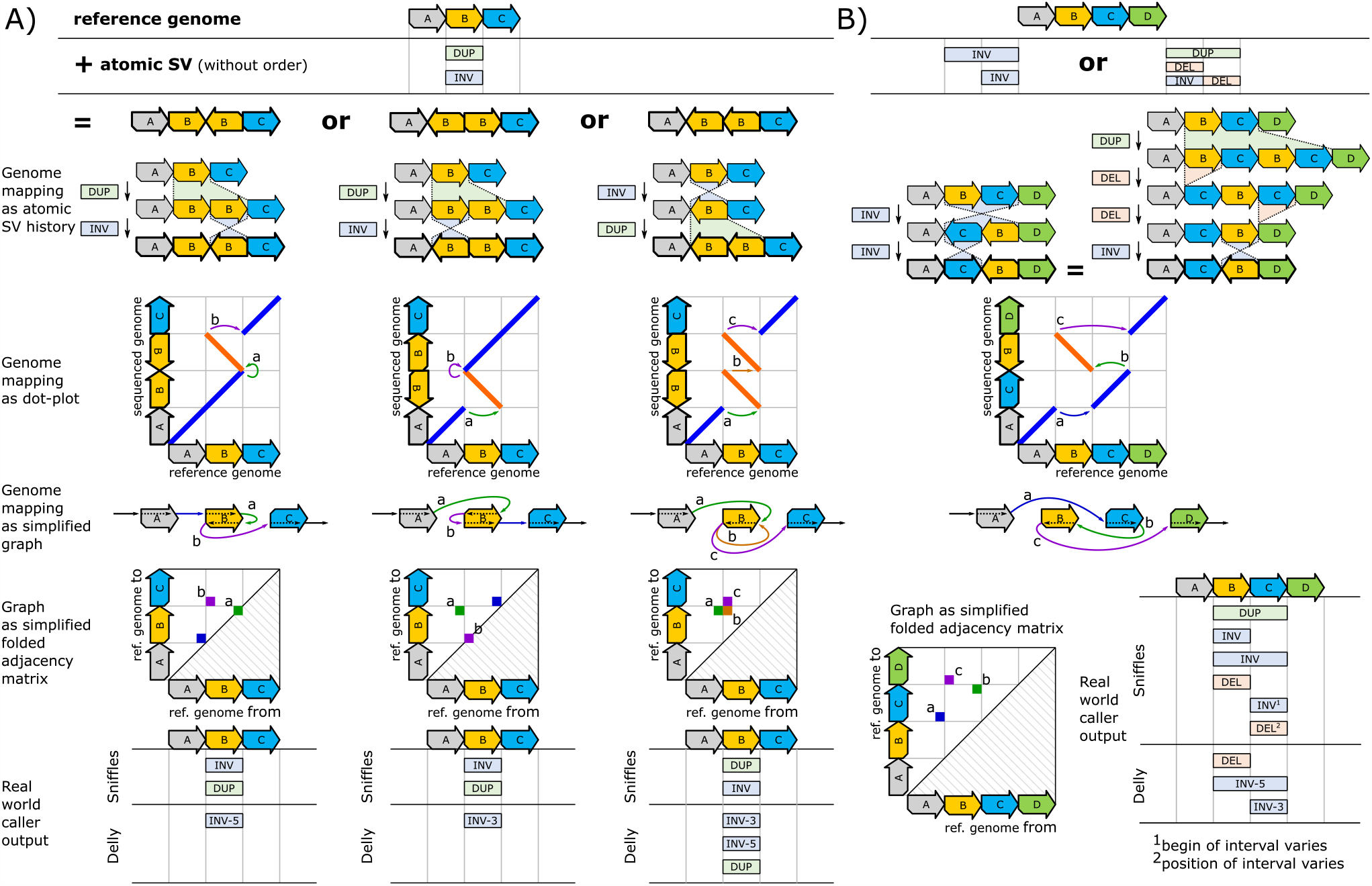
Examples of ambiguities resulting from the description of genomic rearrangements via atomic SV. **A)** shows a situation, where a duplication and an inversion correspond to three different outcomes, which are depicted by diagrammatic dot-plots (for a description of dot-plots and their diagrammatic representation see Supplementary Note 2). Each outcome can be uniquely represented by a graph and its folded adjacency matrix. Here we show simplified versions of the graphs and their folded adjacency matrices merely. The unfolded matrices as well as the full graphs for all examples are given in Supplementary Note 3. The outputs of Sniffles and Delly exemplarily show the behavior of real-world SV callers for the three scenarios. **B)** displays an example, where different rearrangements via atomic SV lead to equal outcomes. Here, a pair of inversions is equal to a duplication followed by two deletions and an inversion. As for A), the corresponding graph, folded adjacency matrix and output of real-world SV callers are shown for both examples. Supplementary Note 3 gives an analogous example, where a duplication followed by two inversions leads to the same outcome as two duplications followed by one inversion.

The abovementioned ambiguities can be eliminated by expressing the outcome of the rearrangement process (instead of the process itself) in terms of the reference. This can be accomplished by e.g. dot-plots as shown in Fig. 1. Each dot-plot, in turn, can be translated into a genome-mapping graph, where all breakend pairs of the dot-plot become edges. (The notion breakend is similar to the notion breakpoint as explained in Supplementary Note 1.) Moreover, the genome-mapping graph can be represented using an adjacency matrix that can be efficiently stored in a database, a CSV-file or by tying together breakends in the VCF-format [27] (using the BND tag and no other tag). For unifying forward and reverse strand, an adjacency matrix can be folded. A detailed definition of our graph-model, the inference of adjacency matrices and their folding scheme is given in the methods section (Fig. 3 and Fig. 6).

**Fig 2.**
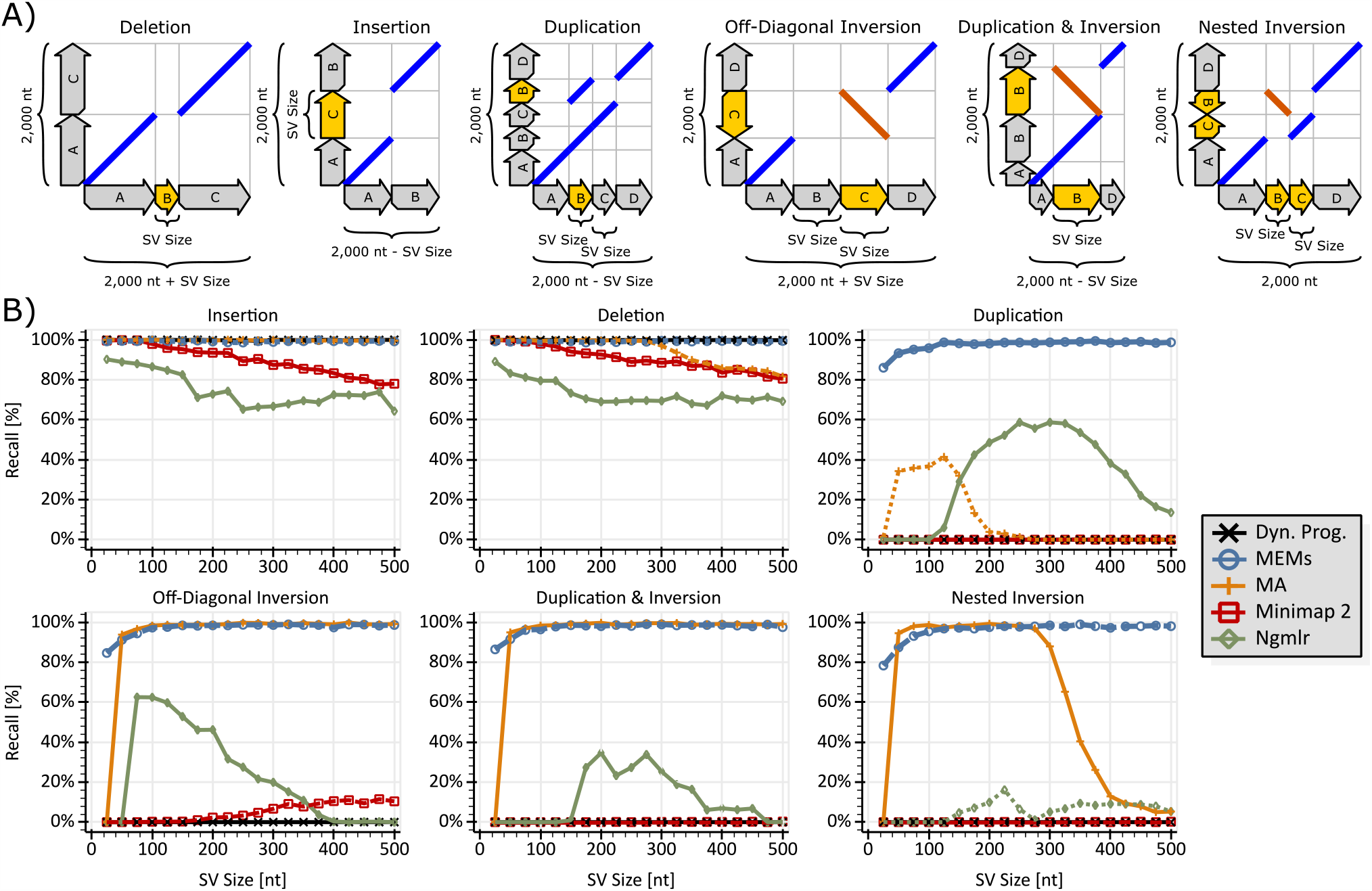
The figure compares MEMs (blue curve), Minimap2 (red curve), NGMLR (green curve) and MA (orange curve) regarding the rediscovery of several types of SV on the foundation of error-free reads. The human genome GRCh38.p12 is used as the reference genome. The black curves (Dyn. Prog.) display the rediscovery rates for banded global Dynamic Programming with two-piece affine gap costs. The global Dynamic Programming is merely applied to the reference-section that a read originates from. Subfigure **A)** visualizes the analyzed genomic rearrangements using diagrammatic dot-plots. Blue lines indicate matches on the same strand, while orange lines represent matches on opposite strands. A description of diagrammatic dot-plots can be found in Supplementary Note 2. The reads for B) are generated by choosing a random section of the reference genome that is rearranged according to the pattern shown in the respective diagrammatic dot-plot of A). Aside from these rearrangements, there are no further modifications (e.g. no simulation of sequencer errors). Subfigure **B)** displays the rediscovery rate as a function of the SV size (see subfigure A) for the various kinds of SV in A). Each point represents an average measurement for 10,000 reads of length 2,000 nt.

**Fig 3.**
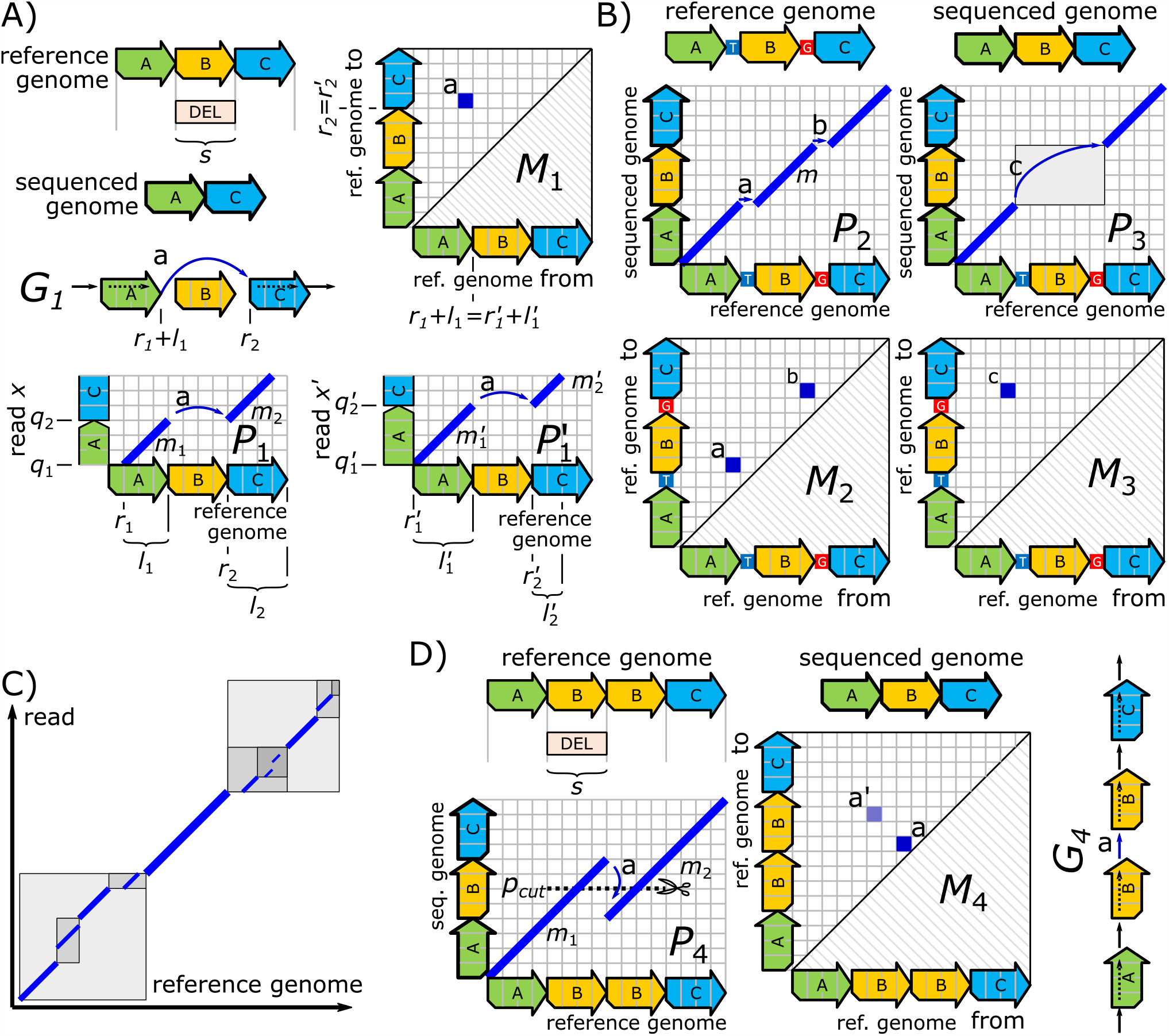
The Figure displays several aspects of our proposed scheme for the computation of genome mappings and their representation using adjacency matrices. **A)** The reference genome’s section *B* is deleted on the sequenced genome. The simplified genome-mapping graph *G*_1_ expresses the sequenced genome in terms of the reference genome, where the edge *a* in *G*_1_ represents the deletion of *B. M*_1_ is the folded adjacency matrix for *G*_1_, where the entry *a* corresponds to the edge *a* in *G*_1_. The entry *a* can be inferred from the MEM’s *m*_1_ and *m*_2_ as well as 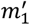 and 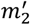 reference positions. These MEMs belong to the read *x* and *x*′, respectively. *P*_1_ and 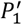 show *a*’s location in form of diagrammatic dot-plots for the reads *x* and *x*′. **B)** Two 1 nt deletions between reference genome and sequenced genome create the small seed *m* in the diagrammatic dot-plot *P*_2_. In *P*_3_, *m* is assumed to be deleted by the mimicked occurrence filtering of our approach (see main text). This deletion leads to the replacement of the two correct entries *a* and *b* in *M*_2_ by the wrong entry *c* in *M*_3_. For salvaging such lost entries, we rediscover *m* via a reseeding within the gray boxed area of *P*_3_. **C)** visualizes our reseeding technique for a larger area, where each box corresponds to a recursive call and the gray’s darkness indicates the recursion depth (darker colors express deeper recursion levels). **D)** shows a problem caused by the repetitiveness of the reference genome. In the sequenced genome, one of the occurrences of *B* is missing. In *P*_4_, the two seeds *m*_1_ and *m*_2_ that overlap on the y-axis, correspond to the erroneous entry *a* in *M*_4_ (and so to the erroneous edge *a* in *G*_4_). Our overlap-elimination technique described in the methods section resolves this problem by shortening *m*_1_ and *m*_2_ to the line *p*_*cut*_. As a result of this shortening, a correct entry *a*′ appears in *M*_4_.

**Fig 4.**
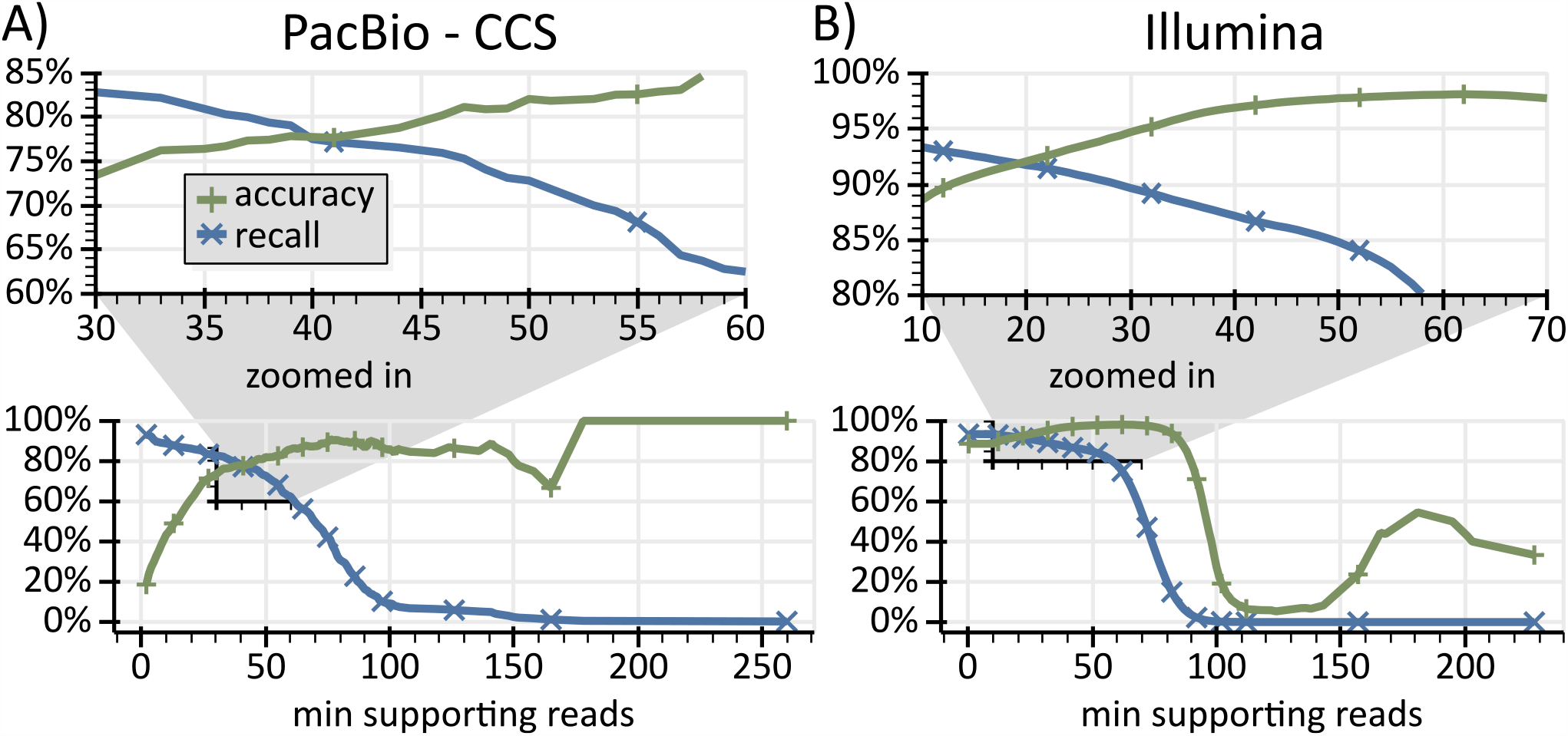
The figure shows exemplary accuracy rates and recall rates of our approach for the two yeast genomes UFRJ50816 and YPS138. **A)** plots two curves that express accuracy rates (green) and recall rates (blue) for matrix entries computed by simulated PacBio reads as functions of an entry’s number of supporting reads. E.g., at x-position 50, the blue curve indicates that for 73% of all ground truth entries there are at least 50 simulated PacBio reads that yield an equivalent matrix entry. Further, the green curve indicates that 82% of the matrix entries supported by 50 (x-position) or more reads are correct (there is a matching ground-truth for these entries). Here a distance of 100nt between ground-truth and matrix entry is granted. **B)** shows accuracy rates (green) and recall rates (blue) for Illumina reads in the same fashion as A) does for PacBio reads. For Illumina reads, the positions of ground-truth and matrix entry must match perfectly.

**Fig 5.**
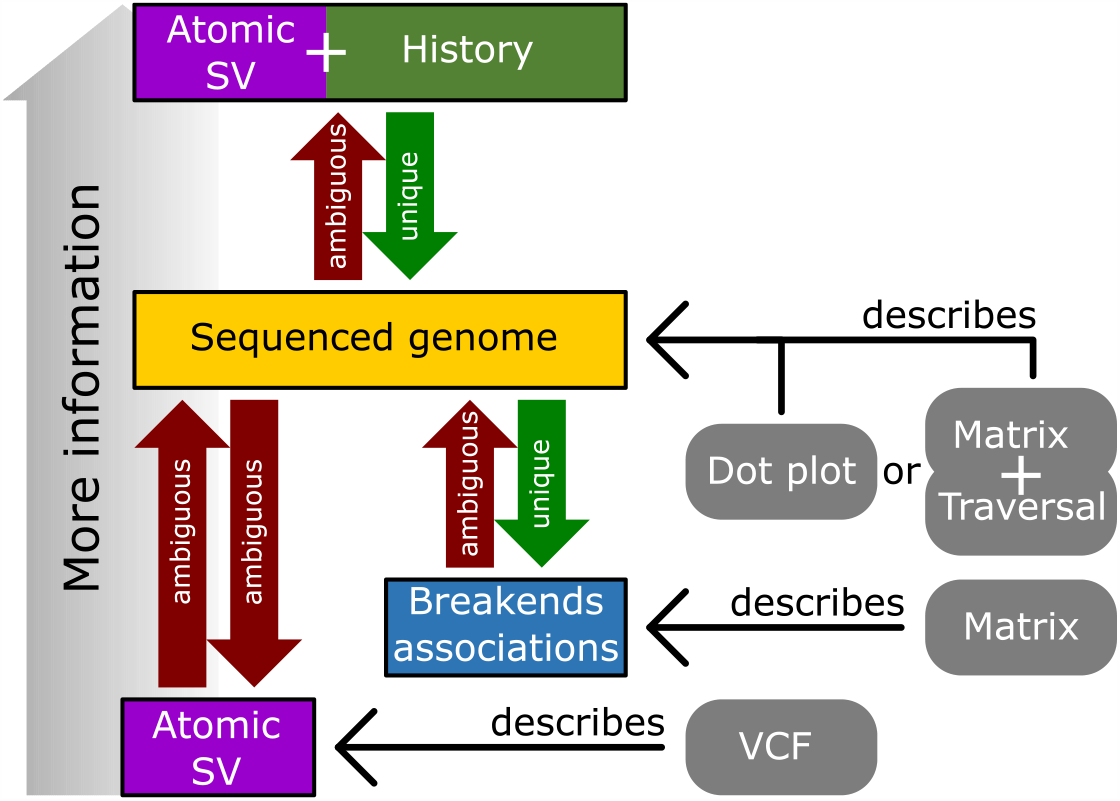
The figure visualizes several ambiguities inherent to the workflow of state-of-the-art SV calling. Further, it places three SV description schemes (dot-plot, VCF-format and adjacency matrix) into their context. Supplementary Note 9 gives examples for the four shown ambiguities (red arrows) together with additional comments.

**Fig 6.**
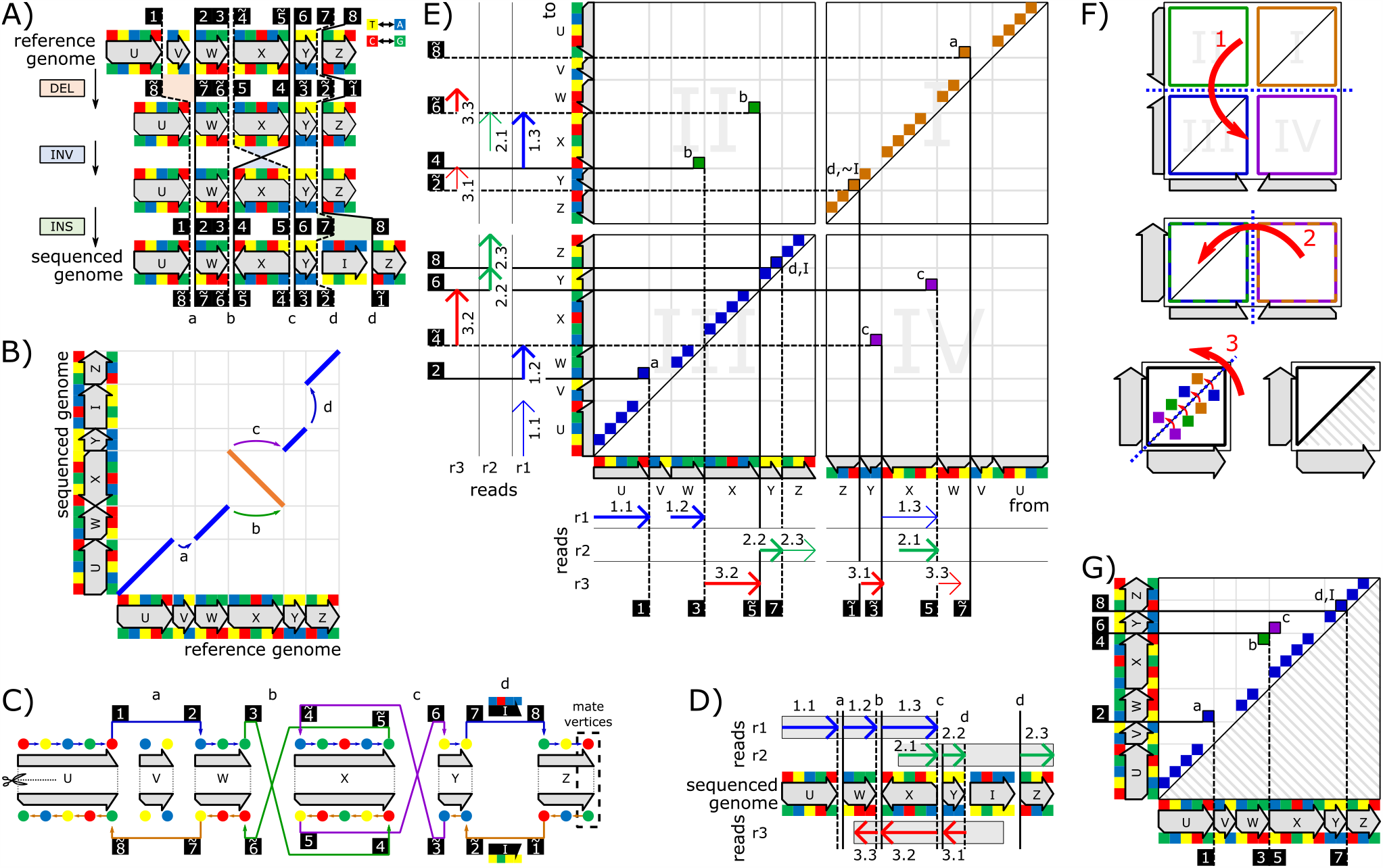
A visual overview of our approach for inferring a folded adjacency matrix from reads. **A)** introduces a reference genome, a sequenced genome and a history of atomic SV (consisting of a deletion of the section *V*, inversion of *X* and insertion of *I*) that transforms the former genome into the latter genome. The black-boxed numbers indicate the order of the breakends on the sequenced genome. A tilde over a number expresses that the corresponding breakend is on the reverse strand of the sequenced genome. The arrowheads of the genome sections *U, V, W, X, Y* and *Z* symbolize their direction on the reference; the colored boxes above and below are their nucleotides (see Supplementary Note 2) on forward and reverse strand, respectively. **B)** shows the genomic rearrangement of A) in form of a diagrammatic dot-plot (details on these dot-plots are in Supplementary Note 2). Each of the breakend pairs *a, b, c* and *d* of A) is indicated via an equally labeled arrow. **C)** displays the skew-symmetric graph for the genomic rearrangement of B). The dashed box on the graph highlights an exemplary pair of mate vertices. The labeled edges of the graph correspond to the equally labeled breakend pairs of A). The weights *I* on the edges labeled *d* represent the inserted sequence on forward and reverse strand. **D)** introduces three error-free reads *r*1, *r*2 and *r*3. Their locations on the sequenced genome are visualized via gray boxes and their MEMs are displayed by colored arrows. *I* is not covered by seeds because it is an insertion. **E)** comprises the unfolded adjacency matrix for the skew-symmetric graph in C). The matrix is inferred from the three reads of D), where the MEMs can be associated via their numbers. For example, the entry *a* corresponds to the two breakends (1) and (2), which are discovered via the MEMs 1.1 and 1.2 of the read *r*1. The first and last seed of each read has no breakend on the y-axis and x-axis, respectively. Such seeds are distinguished by using thin arrows. The edge weights *I* and ∼*I* on the mate edges labeled *d* denote the inserted sequences on forward and reverse strand, respectively. The coloring scheme for the matrix entries memorizes the strand information of edges as described in the “Folding of Adjacency Matrices” chapter of the methods section. **F)** visualizes the adjacency matrix folding scheme of our approach. A step-by-step description of the folding for the matrix in E) is given in Supplementary Note 10. **G)** depicts the folded form of the matrix E). In the folded form, forward and reverse strand are unified. Therefore, all equally labeled entries of the matrix in E) appear as a single entry in G).

We now analyze the practical relevance of the abovementioned ambiguities by inspecting the output of the two popular SV callers Sniffles [28] and Delly [4]. For this purpose, for each dot-plot, we generate a set of perfect alignments, which reflects the rearrangement of the respective dot-plot and fully covers the y-axis. A detailed description of the experimental setup is given in Supplementary Note 4. The generated alignments are forwarded to the SV-callers for evaluation and their output is visualized in the bottom row of Fig. 1. All SV-callers reconstruct the reference positions of the breakends successfully but struggle with the abovementioned ambiguities. For subfigure A), Sniffles reports an inversion and duplication for all three reorganizations. Therefore, Sniffles recognizes the atomic SV correctly but cannot distinguish between the three cases. Delly distinguishes all three cases but does not report the duplication in the first two rearrangements. For Fig. 1 B), Sniffles returns a combination of both development histories, where the atomic SV are sorted by their start position on the reference (due to this sorting both histories appear intermingled). Delly’s output seems to comprise fragments of both development histories. Using our proposed representation scheme, all cases can be represented unambiguously by a unique adjacency matrix.

### Alignments can conceal SV

SV-callers recognize genomic rearrangements using the locations of their breakends. Here many state-of-the-art SV-caller [1, 13, 14] rely on alignments for split-reads (chimeric reads). Therefore, these SV-callers require the aligner to deliver precise and consistent breakend locations. Aligners are designed to find maximally scored paths in the tradition of Needleman-Wunsch’s (NW) [18] and Smith-Waterman’s [19] algorithm. Such a path-based approach is well suited for the discovery of mutations, short insertions and short deletions. However, long insertions and long deletions oppose maximally scored paths. This problem is well known and reflected by the introduction of two-piece affine gap-costs [29]. Further, the path-driven approach contradicts duplications and inversions as indicated by the Dynamic Programming (DP) curves in Fig. 2 B). State-of-the-art aligners attempt to compensate for these shortcomings by identifying their respective cases using tailored strategies and reporting them under the assistance of supplementary alignments. For example, Minimap2 tries to identify potential inversions using a z-drop heuristic during DP and verifies these potential inversions via DP on the reverse complement.

In Fig. 2, we evaluate the aligners Minimap2 [20], NGMLR [28] and MA [21] in the context of the abovementioned shortcomings. There, we benchmark the discovery rates of the aligners for the following SV types: 1) A plain deletion. 2) A plain insertion. 3) A duplication, where a gap occurs between the original and copied segment. 4) A deletion directly followed by an inversion. 5) An inverted duplication (identical to the leftmost case in Fig. 1 A). 6) Two nested inversions (identical to the case in Fig. 1 B2). In all six cases, error-free reads that span over the complete SV are generated by performing the following three steps for each read: 1) We pick a randomly located interval of given size (this size is denoted on the x-axis of the respective case in Fig. 2 A) on the human genome. 2) The sequence in that interval is decomposed into sections as visualized on the x-axis (sections named *A, B, C*, …). 3) An error-free read is generated by copying and rearranging these sections as shown on the y-axis. All reads generated by the above scheme are of equal size (2,000 nt). Following the generation, the reads are forwarded to the aligner for alignment computation. Using the CIGARs of the primary alignments and the supplementary alignments, we verify an aligner’s discovery of all breakends that belong to the respective SV. Here we grant a tolerance of 10 nt and verify reference locations as well as read locations of breakends. An SV counts as rediscovered if all its breakends are found.

Fig. 2 B) indicates the limited success of state-of-the-art aligners in overcoming their path-oriented design in the context of the respective genomic rearrangements. As expected, small deletions and insertions are well discovered by all aligners. However, the discovery rate decreases with increasing indel size. This is in accordance with the abovementioned opposition of the path-oriented scoring scheme to long indels. The DP curves in Fig. 2 B) indicate that two-piece affine gap-costs cope well with larger indels. Therefore, the worse behavior of all aligners for indels is assumed to originate from the seed-processing step (e.g. chaining [30]). The pictures of cases comprising duplications and inversions are mixed, where aligners recognize some cases while failing on others. In contrast, the blue curves in Fig. 2 B) for Maximal Exact Matches (MEMs), where a MEM is a seed that is maximally extended in both directions, illustrate that seeds are sufficient for recognizing all these cases. The failing of aligners can be explained by their path-driven selection of seeds that occurs as part of the seed-processing step. Here aligners purge seeds (MEMs) that are required for recognizing these cases.

Aligners rely on a technique called “occurrence filtering” for coping with the ambiguity of genomes. For keeping fairness, we mimic this occurrence filtering during MEM computation by discarding all MEMs that surpass a given threshold of occurrences on the genome. Except for insertions and deletions, Fig. 2 B) shows a decreasing discovery rate with MEMs for small SV-sizes. This can be explained by the mimicked filtering in combination with the ambiguity of the human genome because small seeds tend to be purged by the occurrence filtering and so the corresponding breakends cannot be discovered anymore. As with the alignments, a breakend is considered as “discovered by a MEM” if one of the MEMs endpoints and the breakend have equal read and reference positions. Further, an SV counts as rediscovered by MEMs if all its breakends are found. Please note that all the cases investigated in Fig. 2 mimic real-world rearrangements that we encounter during our analysis of the yeast genomes UFRJ50816 and YPS138.

Some aligners, as e.g. Minimap2, can be configured to report all discovered chains via supplementary alignments. If configured this way, an aligner’s recall rate in the above analysis increases significantly (as shown in Supplementary Note 5 for Minimap2). However, the configuration results in an abundance of CIGARS without any contextual information among them. Without this contextual information, the CIGARs are of similar value as pure MEMs in the context of SV calling.

### SV calling using genome-mapping graphs inferred from MEMs

The blue curves in Fig. 2 B) suggest that SV calling using MEMs is advantageous. We now outline our technique for computing genome-mapping graphs using MEMs. The technique is visualized in Fig. 3 A). In the following, we assume that we have an error-free read *x* and two consecutive MEMs *m*_1_ = (*q*_1_, *r*_1_, *l*_1_) and *m*_2_ = (*q*_2_, *r*_2_, *l*_2_) for *x*. Here, the *q*’s and *r*’s are the start positions of the MEMs on the read and reference, respectively. *l*_1_ and *l*_2_ are the lengths of the MEMs. The consecutive MEMs *m*_1_ and *m*_2_ indicate a difference between the read *x* and the reference. For example, if this difference is a deletion of size *s*, then *m*_1_ and *m*_2_ touch each other on the read *x* (because *q*_2_ = *q*_1_ + *l*_1_) but there is a gap of size *s* between them on the reference (*r*_2_ = *r*_1_ + *l*_1_ + *s*). We transfer this difference information into the adjacency matrix of the genome-mapping graph by creating an entry (*r*_1_ + *l*_1_, *r*_2_) in the matrix. Informally, by doing so, we store the information that the read comprises a “jump” from *r*_1_ + *l*_1_ to *r*_2_ on the reference. By relying on such a scheme, we can uniquely store arbitrary differences between reads and reference genome. Further, all reads that span over a difference confirm the same matrix entry because the read positions of MEMs are not incorporated into the matrix entries. The adjacency matrix, in turn, uniquely defines the genome-mapping graph (the detailed scheme is described in the methods section). Such an adjacency matrix must store information regarding forward strand and reverse strand. In the methods section, we propose a matrix-folding scheme that unifies the entries of forward and reverse strand reads. Due to this folding scheme, reads of both strands can indicate the same rearrangement. A detailed description of our SV calling approach is given in the methods section. This includes the proposal of a memory-efficient representation of our adjacency matrices.

### Incorporating genome repetitiveness and sequencer errors into SV calling

Repeats on the reference genome can cause MEMs to overlap on the y-axis (on the read) as shown in Fig. 3 D). There, the MEMs *m*_1_ and *m*_2_ both cover the section labeled *B* on the y-axis. This overlap leads to the erroneous matrix entry labeled *a*. This entry expresses that, on the sequenced genome, the reference’s first instance of *B* is followed by the reference’s second instance of *B*. However, for correctly representing the sequenced genome via the adjacency matrix, we instead have to express that there exists only one instance of *B* on the sequenced genome. A graph genome, where such repetitiveness is modeled by edges instead of vertices, would eliminate this problem (more details on graph genomes can be found in the discussion section). In the methods section, we introduce an algorithmic scheme that shortens overlapping seeds for resolving such overlaps on sequential reference genomes.

The repetitiveness of genomes creates further issues. In the following, let *S* be a sequence that occurs once on the reference genome but *n* times on the sequenced genome. This leads to a situation, where the vertex belonging to the first or last nucleotide of *S* comprises *n* inbound or outbound edges, respectively. Relations between such inbound and outbound edges cannot be expressed by the graph itself. Instead, they must be defined by a graph traversal. Since our approach currently does not compute such a traversal, we avoid these regions by applying a filtering approach on all reads as described in the methods section. Now, let *S* be a sequence that occurs *n* times on the reference genome and at least once on the sequenced genome. In this case, an occurrence of *S* on a read triggers *n* matches on the reference, where all of them are false positives except for one. For handling this situation, we discard all seeds. This technique is similar to the “occurrence filtering” that is part of the seeding step of many aligners [20, 21, 31]. The discarding of ambiguous seeds primarily removes small seeds since the number of occurrences of a seed on a genome is expected to increase with decreasing size of the seed. However, small seeds are required if two differences between read and reference are close to each other. Such differences can be caused by genomic rearrangements, sequencer errors or combinations of them. If the seed between two differences is missing, two correct entries in the adjacency matrix are replaced by a single wrong entry (see Fig. 3 B). For tackling this kind of problem, we propose an efficient reseeding scheme for the retrieval of small seeds on a locally confined region of the reference (see Fig. 3 C). The required basic techniques for obtaining such seeds using an index computed on the fly are proposed in [32]. There, a hash table is used for computing Minimizers that are merged and extended afterward for obtaining MEMs. For each MEM, we search for smaller undiscovered MEMs occurring immediately before and after it. This process is recursively extended to freshly discovered MEMs until they are expected to be too small (down to 5nt). The detailed process is described in the Methods section.

### Evaluation using yeast genomes

So far, we analyzed individual aspects of our SV calling approach merely. Now, we perform a comprehensive evaluation using the real-world data proposed in [22] for *Saccharomyces paradoxus* (wild yeast). We work with yeast genomes due to their small size and moderate repetitiveness in comparison to e.g. the human genome. In the Discussion section, we outline extensions and modifications of our approach that would allow its application to the human genome. The data in [22] are particularly well suited for our analysis because they comprise independently assembled genomes for several strains of wild and domestic yeast, which can be used as ground truth for our analysis. We focus on the genomes UFRJ50816 and YPS138 here. These genomes comprise complex genomic rearrangements (e.g. six clustered inversions, some of which are overlapping) that are similar to the situations shown in Figure 1 B2. Finally, the yeast genomes in [22] are all sequenced in their haploid or homozygous diploid forms, which simplifies SV calling compared to diploid genomes comprising heterozygous loci.

Due to the reported ambiguities of atomic SV, an evaluation of our approach using VCF-formatted data as ground truth is not feasible. For overcoming this problem, we compute a set of adjacency matrix entries based on a reference assembly *A*_*R*_ (we use YPS138) and a sequenced assembly *A*_*S*_ (we use UFRJ50816) as ground truth entries. The entries are inferred from a comprehensive set of MEMs for *A*_*R*_ and *A*_*S*_. Here, we rely on the same algorithmic techniques for computing seeds and matrix entries as we use for reads (see methods section). The overall scheme for the computation of the ground-truth entries is given in Supplementary Note 6. For the ∼12Mio bp long yeast genomes, this ground-truth matrix denoted as *M*_*T*_, comprises 43,154 entries.

The theoretic foundations of our approach demand that *A*_*S*_ corresponds to one specific traversal throughout the genome-mapping graph, where the graph is defined by the ground-truth entries and the traversal is determined by a visiting order among the entries. For UFRJ50816 (as *A*_*S*_) and YPS138 (as *A*_*R*_) we could fully reconstruct the sequenced genome using a prototype implementation, which is described in Supplementary Note 7. This, in turn, proves the correctness of the practically computed ground-truth entries for our pair of *A*_*S*_ and *A*_*R*_.

So far, for the generation of the ground-truth entries, the complete genome *A*_*S*_ is used as one virtual error-free read. Now, we evaluate our approach using simulated reads, where we rely on SURVIOR [33] and DWGSIM [34] for the generation of CCS-PacBio and Illumina reads, respectively. The error profiles for SURVIOR’s CCS-PacBio read generation are sampled from Minimap2 alignments for the PacBio-MtSinai-NIST reads of HG002 (AJ Son) [35]. Let *M*_*PacBio*_ and *M*_*Illumina*_ be the two matrices computed for the simulated PacBio and Illumina reads. PacBio reads are well suited for the detection of larger genomic rearrangements, while Illumina reads are appropriate for the discovery of short rearrangements. Therefore, we split the 43,154 entries in *M*_*T*_ into two submatrices: A matrix *M*_*T,large*_ consisting of the 320 entries in *M*_*T*_ that connect breakends further apart than 200nt on the sequenced genome or reference genome. All entries of *M*_*T*_ that are not in *M*_*T,large*_ form a submatrix *M*_*T,small*_. The matrix *M*_*T,large*_ is used as ground-truth for the analysis of *M*_*PacBio*_ that is shown in Fig. 4 A). Accordingly, Fig. 4 B) displays the outcome of the analysis for *M*_*Illumina*_ with *M*_*T,small*_ as ground-truth. In both subfigures, the recall rate drops-off starting at ∼50 reads (x-axis), which indicates that for almost all rediscovered entries more than half of the supporting reads are found by our approach. Sequencer errors and small variations (e.g. SNP or small indels) in repetitive regions can create false positives. However, most of these false positives can be distinguished from true positives because they are supported by less than ∼ 30 reads. By using simulated reads instead of the real-world reads provided by [22], we guarantee that potential errors of the assembly process do not affect our analysis.

The outcome of SV calling should allow the reconstruction of the sequenced genome from the respective reference genome. Our genome mapping graph model (adjacency matrices) permits such a reconstruction as described in the methods section. We now evaluate the genomes that result from a reconstruction using *M*_*T*_ as well as *M*_*PacBio*_ ∪ *M*_*Illumina*_. As shown in Table 1, *M*_*T*_ delivers a perfect reconstruction of the sequenced genome, which justifies the use of *M*_*T*_ as ground-truth. With *M*_*PacBio*_ ∪ *M*_*Illumina*_, the identity between chromosomes of the reconstructed genome and the sequenced genome varies from 97.0% to 99.9%. Here, these identities are computed as 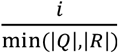, where *i* represents the number of matches in a banded (bandwidth = *abs*(|*Q*| − |*R*|) + 10,000nt) Needleman Wunsch alignment with two-piece affine gap costs between the sequences *Q* and *R*. As described in the methods section, the reconstruction requires a traversal. For now, we do not compute this traversal for *M*_*PacBio*_ ∪ *M*_*Illumina*_. Instead, we use the traversal of *M*_*T*_ as an approximation. However, this approximation can contradict the graph defined by *M*_*PacBio*_ ∪ *M*_*Illumina*_. Such contradictions occur between 226 (0.6%) of the 40,270 entries in *M*_*PacBio*_ ∪ *M*_*Illumina*_. The approximation could be replaced by a naïve traversal that always prioritizes non-implicit edges over implicit edges (see the first subsection of the methods section for the definition of implicit edges) and stops if multiple non-implicit edges originate from a vertex. 94.3% of pairs of consecutive edges in the approximated traversal are part of such a naïve traversal as well, i.e. merely 5.7% of edge pairs of the approximation are contributing. Comparing the naïve approach to the traversal of *M*_*T*_ on *M*_*T*_’s graph itself, 97.6% of pairs of consecutive edges are part of both traversals. 4.5% and 3.1% of the reconstruction via *M*_*T*_ and *M*_*PacBio*_ ∪ *M*_*Illumina*_ originate from the weights (inserted sequences) of entries, respectively. 42,597 entries in *M*_*T*_ and 37,726 entries in *M*_*PacBio*_ ∪ *M*_*Illumina*_ have weights. This use of weights is necessary since some differences between the reference genome and the sequenced genome (e.g. mutations and insertions) should not be expressed via sequences of the reference genome. In our case, sequences that are not covered by any seed are represented as weights. This implies that some duplications become weights due to the occurrence filtering (see methods section). Here, 90.6% and 93.8% of these weights are sequences of size one (mutations), while 0.6% and 0.5% of these weights are sequences of size ≥ 100 nt (long insertions or long repetitive regions), for *M*_*T*_ and *M*_*PacBio*_ ∪ *M*_*Illumina*_, respectively. Large insertions connect to one of two special sentinel vertices as described in the methods section. However, in the context of our evaluation of Yeast, all insertions are fully enclosed by one PacBio read at least. Therefore, for all entries, the merging scheme described in Supplementary Note 8 can be applied in the context of the entries’ scoring.

**Table 1.** The table displays the rates of identity between the sequenced genome and its reconstructed forms using *M*_*T*_ and *M*_*PacBio*_ ∪ *M*_*Illumina*_. For comparison purposes, the identity between reference genome (*A*_*R*_) and sequenced genome is shown as well. The scheme used for calculating the identity is described in the main text.

|  |  | Chromosome | 1 | 2 | 3 | 4 | 5 | 6 | 7 | 8 |
| --- | --- | --- | --- | --- | --- | --- | --- | --- | --- | --- |
| Identity to<br>Sequenced<br>Genome | | $M_T$ | 100% | 100% | 100% | 100% | 100% | 100% | 100% | 100% |
| | | $M_{PacBio} \cup M_{Illumina}$ | 99.1% | 99.3% | 99.2% | 99.9% | 99.8% | 97.1% | 99.9% | 98.8% |
| | | $A_R$ | 97.4% | 85.8% | 98.5% | 72.5% | 92.3% | 97.8% | 94.5% | 97.7% |
|  |  | Chromosome | 9 | 10 | 11 | 12 | 13 | 14 | 15 | 16 |
| Identity to<br>Sequenced<br>Genome | | $M_T$ | 100% | 100% | 100% | 100% | 100% | 100% | 100% | 100% |
| | | $M_{PacBio} \cup M_{Illumina}$ | 98.9% | 98.6% | 99.7% | 99.9% | 99.8% | 99.7% | 99.7% | 99.9% |
| | | $A_R$ | 76.1% | 97.6% | 83.9% | 90.6% | 85.5% | 88.3% | 93.8% | 91.6% |

## Discussion

In the results section, we investigate ambiguities resulting from the description of genomic rearrangements using atomic SV (e.g. descriptions via the VCF-format). Fig. 5 extends this inspection of ambiguities to the relationships between genomic rearrangements, atomic SV, sequenced genomes and breakend associations: Evolutionary development is a stepwise process; its genetic aspects are usually observed via sequenced genomes, which are snapshots of a specimen’s genetic definition. Aside from some trivial cases, this snapshot-approach cannot reveal the history of genomic alterations (atomic SV) that turned one snapshot into another because different histories can lead to the same outcome. For example, three inversions can alter a sequence in the same way as one duplication followed by two deletions as e.g. shown in Supplementary Note 9 (subfigure D). This inherent ambiguity, in turn, affects all atomic SV-based description schemes as e.g. the VCF-format. Therefore, any quantitative measurement of atomic SV has to exclude nested variants (e.g. by using the benchmarking dataset proposed in [23]) or it is doomed to be ambiguous. For example, in [24-26] these ambiguities are tackled by defining signature variant allele structures for several complex genomic rearrangements while ignoring all others. Further, the repetitiveness of genomes in combination with the limited length of sequenced reads inflicts an additional level of ambiguity on the snapshots (sequenced genomes) themselves. For example, if a duplication’s size exceeds the length of all reads, it is impossible to reconstruct its number of repeats (see Supplementary Note 9, subfigure E). Our adjacency matrix-based approach resolves these ambiguities in two ways: 1) It abstracts from the history of genomic alterations. 2) It differentiates between the adjacency matrix itself and a traversal through its respective graph. In this context, the adjacency matrix captures the locations of all breakend pairs (see Supplementary Note 1 and Fig. 6), while the traversal corresponds to the occurrence order of the breakend pairs on the sequenced genome. This traversal can only be computed if there are reads of sufficient length. However, for inferring adjacency matrix entries, short reads are sufficient since each entry merely requires two seeds to identify its breakend pair.

The repetitiveness of genomes generally poses a problem during the analysis of genomic data. In this context, the repetitiveness of sequenced genomes affects our approach differently than the repetitiveness of reference genomes. A sequence that occurs once on the reference genome but many times on the sequenced genome equals one or several duplications. Such duplications create cycles in our graph model that have to be resolved via a graph traversal. On the contrary, sections of a sequenced genome that match several locations on a reference genome are more challenging. Here it is necessary to select the correct location from many candidates on the reference. This selection can be done via alignments; however, as reported in the results section, alignments can conceal nested SV. Instead of tackling the repetitiveness of a reference genome, it would be advantageous to avoid such repetitiveness in the first place. This can be achieved by switching to graph genomes, where repetitive sequences appear only once and the repetitiveness is represented by a path throughout the graph. Currently, the axes of our adjacency matrices correspond to sequential genomes and therefore all matrix entries that connect non-neighboring vertices (vertices that belong to non-neighboring nucleotides on a genome) represent SVs. However, our approach does not require such sequential genomes in the context of the adjacency matrix. Instead, we can use a graph genome as a reference for SV calling by placing all sequences that are part of the graph genome on the axes. Compared to a sequential genome, the genomic rearrangements are now matrix entries that connect vertices not neighboring in the graph genome. Further, the graph traversal that is computed for representing the sequenced genome can immediately be used as another path in the graph genome. Hence, our approach can be used for adding new specimens to pan-genomes, since pan-genomes are usually considered to be graph genomes. Such additions might imply a modification of the pan-genome’s graph if the sequenced genome indicates an additional level of repetitiveness. In this work, we use sequential reference genomes for two reasons: 1) Currently, we do not compute the traversal that delivers a sequenced genome. 2) A bootstrapping is required for providing repetition-free graph genomes as starting points.

The yeast genomes used for our analyses are small compared to the genomes of e.g. mammals. This raises the question of the applicability of our approach to larger, more complex and diploid genomes as e.g. the human genome. Diploid genomes do not affect the computation of the adjacency matrix. They merely require a separate traversal for each chromosome of a pair. Currently, the repetitiveness of complex genomes practically overloads the seed processing involved in our approach (SoC, reseeding and overlap elimination). The availability of a repetition-free graph genome would immediately remove this overload. As an intermediate solution for large genomes, our approach could be used on the foundation of alignments, where CIGARs take the position of seeds in the context of the matrix entry computation. However, this would conceal many complex genomic rearrangements as mentioned above.

## Methods

### A skew-symmetric graph model for describing genomic rearrangements

We describe rearrangements between a reference genome *R* and a sequenced genome *S* using a weighted, directed skew-symmetric graph (skew-symmetric graphs are defined e.g. in [36]). In our graph model, vertices represent nucleotides of *R*. Using the edges and their weights, we express *S* in terms of *R*. Let *u* and *v* be two vertices. Further, let *N*(*u*) and *N*(*v*) be their respective nucleotides: An edge from *u* to *v*, indicates that *N*(*v*) follows *N*(*u*) on *S*. Hence, edges can either express equality between *S* and *R*, by connecting the vertices of nucleotides that follow each other on *R*, or express differences (i.e. breakend pairs) by connecting the vertices of nucleotides that are nonconsecutive on *R*. For example, in Fig. 6 C), the edge between the first two vertices of the segment *U* on the forward strand indicates that the corresponding nucleotides appear consecutively on the sequenced genome. Accordingly, the edge labeled *a* expresses that *W* follows *U* on the sequenced genome. For each position on the reference, we have one vertex on the forward strand and one vertex on the reverse strand. These vertices are called mates. Additionally, each edge (*u, v*) has a mate edge that connects the mate vertices of *u* and *v* in opposite direction. In Fig. 6 C), the forward-strand edge labeled *a* has a mate on the reverse strand that connects the first vertex of *W* with the last vertex of *U*. The concept of mates is part of the skew-symmetry of our graph model. Using strand-crossing edges, we can represent breakend pairs of inversions. For example, in Fig. 6 C), the first breakend pair of the inversion of *X* on the sequenced genome is represented by the green mate edges labeled *b*. Insertions are modeled using the weights of edges. Hence, the edge connecting *Y* and *Z* in Fig. 6 C) comprises *I* as weight for modelling the insertion of the sequence *I* on the forward strand. For representing a genome, it is necessary to have a set of traversals that accumulatively visit all non-isolated vertices of the graph at least once. Each traversal represents one chromosome of the sequenced genome.

The example reads of Fig. 6 D), E) demonstrate the inference of entries for an unfolded adjacency matrix in the idealized situation of error-free reads. For this purpose, we use Maximal Exact Matches (MEMs). MEMs are seeds that are maximally extended in either direction. Each entry is generated by the combination of the head (x-position of entry) and tail (y-position of entry) of two arrows, which correspond to the endpoints of two MEMs that occur consecutively on the respective read. For example, the MEMs labeled 1.1 and 1.2 in Fig. 6 D), E) create the entry labeled *a* in quadrant *III*. If a read from the forward strand and a read from the reverse strand span over the same breakend pair, they create mate entries in the unfolded matrix. For example, the mate entries labeled *b* result from read *r*_1_ (on the forward strand) and read *r*_3_ (on the reverse strand). The entries touching the diagonal on their bottom right corner can be directly inferred from the seeds, where such a matrix entry (*u, v*) is formed by the pair of consecutive nucleotides *N*(*u*), *N*(*v*) in the respective seed. We call these entries implicit because they are not related to SVs and can be neglected in practical implementations.

### Folding adjacency matrices

We now propose a folding scheme for unifying mate matrix entries (e.g. the entries labeled *b* in Fig. 6 E) by exploiting the skew-symmetry of our graph model. In the context of this folding, for each edge (*u, v*), we store the strands of *u* and *v* via two annotations. Each annotation can receive the value *F* or *R* for forward strand or reverse strand, respectively. In short, we write *FF, FR, RF* or *RR* for the strand annotations of both vertices of an edge. In the context of these annotations, we use the following coloring scheme: *FF* = blue, *FR* = green, *RF* = purple, *RR* = light brown. In the unfolded matrix, these annotations are intrinsically defined via the quadrant an edge appears in: Quadrant *I* → *RR*, Quadrant *II* → *FR*, Quadrant *III* → *FF*, Quadrant *IV* → *RF* (see Fig. 6 E). The annotations allow a reconstruction of the unfolded matrix out of the folded one.

The folding scheme is visualized in Fig. 6 F) and comprises the following three steps:

1. The top half of the matrix is mirrored to the bottom half of the matrix.
2. The right half of the resulting rectangular matrix is mirrored to the left half.
3. In the resulting matrix, all entries below the diagonal are mirrored to their corresponding mates above the diagonal. During this mirroring, the two binary annotations of entries are swapped and complemented (*FF* → *RR, FR* → *FR, RF* → *RF, RR* → *FF*). The weights of mirrored edges are reverse complemented. Entries on the diagonal that are annotated with *FF* stay untouched, while all other entries on the diagonal are mirrored to themselves as described above.

In Fig. 6 the example matrix of subfigure E) is folded into the matrix shown in subfigure G). The two breakpoints (two breakend pairs) of an inversion always correspond to two entries in the folded matrix, where one entry is annotated *FR* and the other *RF*. For example, in subfigure G), the entries labeled *b* and *c* correspond to the inversion *X*. For isolated inversions, these two entries are always direct neighbors in the folded matrix and their distance to the diagonal indicates the size of the inversion. The deletion of *V* is represented by the mate entries labeled *a*. In the unfolded matrix, one of these mates is annotated *FF* and the other *RR*. The folding combines these two mates as shown in subfigure G). As with isolated inversions, the distance to the diagonal in the folded matrix indicates the size of isolated deletions. The insertion of *I* is represented via the weights *I* and Ĩ (the reverse complement of *I*) on the mate edges labeled *d*. Once more, the matrix folding combines both mates.

### Computing a folded adjacency matrix from real-world reads

In the following, we infer an adjacency matrix from a given set of reads and a reference genome *R*. Initially, we compute the MEMs of all reads. Each pair of these MEMs that occur consecutively on a read creates a single entry in the adjacency matrix. Here multiple reads can create the same entry. For coping with this, we move to a multigraph model. Further, the endpoints of MEMs can deviate slightly from actual breakends due to sequencing errors and the repetitiveness of genomes. Therefore, we include a concept of fuzziness by extending each matrix entry to an area around it. For finding true breakend pairs, we merge overlapping areas and later reduce them to single points. In the context of this merging, the size of these areas should be chosen so that we do not merge areas that belong to different breakend pairs. For dealing with large insertions that are not enclosed by any read, two sentinel vertices are added to our graph model.

### Computing MEMs for a read via reseeding

The MEMs used for inferring the adjacency matrix are computed via a process that comprises three steps:

1) An initial computation of MEMs with respect to the whole reference *R* and a given read, 2) a locally confined reseeding and 3) the elimination of overlaps between MEMs on the same read.

For 1), we compute MEMs of a specific minimum size using Minimizers [37] and a merge-extend scheme [32]. In this context, we apply an occurrence filtering that eliminates hash table entries of Minimizers with too many matches on the reference genome or the sequenced genome. (This is required for handling the repetitiveness of the involved genomes. Please note that the sequenced genome is not directly available. Instead, we use all reads, which incorporate the sequenced genome.) In contrast to the commonly used occurrence filtering on the reference, the occurrence filtering on the sequenced genome is a novel addition. It is achieved in two steps: First, the number of occurrences is counted for all Minimizers on all reads. Next, we compute the reference positions for all minimizers with fewer occurrences than a given threshold, where this threshold is modulated by the average coverage of all reads.

2) Matches smaller than the minimal size of Minimizers as well as matches for repetitive sections purged by the occurrence filtering are missing in the set of MEMs computed in 1). Many of these missing MEMs can be discovered via a locally confined reseeding that works as follows: First, we sort each read’s MEMs according to their start positions on the read. Afterward, we visit both endpoints of each MEM and search in a rectangular area that starts on the endpoint and extends away from the MEM on read and reference (see Fig. 3 C)). If a MEM’s successor or predecessor (on the read) extends into this rectangular area, we shrink the area so that it lies between the endpoints of both MEMs. The size of this area determines the minimal size of MEMs to be discovered by the reseeding using the following formula:

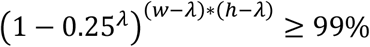

Let *λ* denote the minimal size of MEMs. *w* and *h* are the width and height of the rectangular area for reseeding, respectively. *λ* is chosen so that the probability of a random match of size *λ* in an area of size *w* ∗ *h* is lower than 1%. For each rectangular area, we reseed by computing a single-use Minimizer index and by utilizing the merge-extend scheme proposed in [32] for turning the Minimizers into MEMs. The above procedure is repeatedly applied until it does not deliver new MEMs.

3) Both of the above steps can deliver MEMs that overlap on a read. (These MEMs must originate from the same read.) However, as shown in the results section, such overlapping MEMs lead to erroneous matrix entries. We solve this problem by applying a seed shortening, where we distinguish between two kinds of situations as follows: A) A MEM *m*_1_ is fully enclosed by another MEM *m*_2_ on the read. In this case *m*_1_ is deleted and *m*_2_ is kept unaltered. B) The endpoint of *m*_1_ is above the start point of *m*_2_, but *m*_1_ is not fully enclosed by *m*_2_. In this case, we determine the central point *p*_*cut*_ of the overlap between both seeds and shorten both seeds with respect to *p*_*cut*_. (The endpoint of *m*_1_ is moved to *p*_*cut*_ − 1; the startpoint of *m*_2_ is moved to *p*_*cut*_.) The described overlap elimination is applied on SoCs as well. Supplementary Note 11 visualizes both forms of overlap elimination and extends the explanation.

### Fuzzy inference of edges from MEMs

Our fuzziness concept exploits the spatial locality of neighboring vertices on the reference genome. (Neighboring vertices (and so adjacency matrix entries) correspond to neighboring nucleotides on the reference.) In the following, we represent a MEM as a triple (*q, r, l*), where *q* is the MEM’s start on the read (query), *r* is the MEM’s start on the reference and *l* is the length of the MEM. Let *m*_1_ = (*q*_1_, *r*_1_, *l*_1_) and *m*_2_ = (*q*_2_, *r*_2_, *l*_2_) be two MEMs that occur consecutively on some read. These two MEMs create a matrix entry *e* = (*v*_*x*_, *v*_*y*_) with *x* = *r*_1_ + *l*_1_ and *y* = *r*_2_. Here, the vertex *v*_*i*_ corresponds to the *i* th nucleotide of the reference genome. Further, we assume the existence of a corresponding true entry, denoted by *T*(*e*), which emerges from error-free reads. The entry *e* can deviate from *T*(*e*) due to sequencing errors that shorten or extend MEMs erroneously. Here a shortened MEM causes a deviance leftwards (*m*_1_ shortened) or upwards (*m*_2_ shortened) and an enlarged MEM causes deviance rightwards (*m*_1_ extended) or downwards (*m*_2_ extended). An erroneous extension of *n* nucleotides requires a wrongful match between reference and read of *n* nucleotides, which is caused by *n* consecutive sequencer errors. On the contrary, a shorting by *n* nucleotides can be caused by a single sequencer error that breaks a MEM into two pieces, where the small piece (required for the entry) gets lost due to size constraints or occurrence filtering. This implies that an erroneous shortening is more probable than an erroneous enlargement. For dealing with such deviations of entries, we define a rectangular area around each entry *e* = (*v*_*x*_, *v*_*y*_) that is expected to include *T*(*e*) as follows: For denoting a rectangular matrix area *A*, we use a quadruple (*a, b, w, h*), where *a, b* are the coordinates of the top-left point of *A* and *a* + *w, b* − *h* are the coordinates of the bottom-right point of *A*. We define a rectangular area *A*(*e*) = (*x* − *σ, y* − *σ, f* + *σ, f* + *σ*), where *f* and *σ* deal with the erroneous shortening or enlargement of MEMs, respectively. *A*(*e*) is called the entry area of *e* and we choose *f* = min(|*x* − *y*|, 10) as well as *σ* = 3 by default. Supplementary Note 12 visualizes the concept of entry areas. All entry areas *A*(*e*) inherit a strand annotation from their respective entry *e*. The folding of the adjacency matrix automatically mirrors entry areas and adapts their strand annotations as required. Accordingly, in the folded matrix, an entry area can extend into four different directions with respect to its origin. For coping with the high density of entries immediately above the diagonal of the adjacency matrix, the fuzziness parameter *f* is downregulated in this region (see Supplementary Note 13). For an adjacency matrix *M*, entry areas in *M* correspond to bipartite subgraphs of *M*’s graph.

### Computing clusters by merging overlapping entry-areas

Let *M* be a matrix and *E* be the maximal set, where we have *T*(*e*) = *T*(*e*^′^) for all pairs *e, e*^′^ ∈ *E*. This entry, which is equal for all elements in *E*, is called the true entry of *E* and denote it by *T*(*E*). If the parameter *f* and *σ* are properly chosen, the entry area *A*(*e*) overlaps *T*(*E*) for all entries *e* ∈ *E*. This implies that all these entry areas mutually overlap. Hence, we can compute the set *E* (without knowledge about the true entry *T*(*E*) itself) by joining overlapping entry areas of *M* into clusters. Such clustering can be performed efficiently by a single line-sweep as described in Supplementary Note 13. Supplementary Note 12 B) shows an example for the clustering of several entry areas that belong to the same true entry. As described above, the matrix folding maps areas originating from the reverse strand into the corresponding areas of the forward strand. The clustering is performed after the folding so that entry-areas from forward and reverse strand reads are unified. The weights (inserted sequences) of matrix entries in *E* can be different. In this case, multiple sequence alignments can be used for obtaining a unified weight. So far, we skip such multiple sequence alignments and leave the weight empty instead.

### Approximating true entry locations

We now use all entries in *E* for computing an approximation of *E*’s true entry *T*(*E*). For this purpose, we compute two sets *X* and *Y* as follows: *X* = {*i* | (*v*_*i*_, _) ∈ *E*}, *Y* = {*j* |(_, *v*_*j*_) ∈ *E*}. Depending on the two strand-annotations of *E* (these must be equal for all *e* ∈ *E*), the approximation of *T*(*E*) = (*u*_*x*_, *u*_*y*_) is defined as follows:

*FF*: *x* is the 95^*th*^ percentile of *X* and *y* is the 5^*th*^ percentile of *Y*

*FR*: *x* is the 95^*th*^ percentile of *X* and *y* is the 95^*th*^ percentile of *Y*

*RF*: *x* is the 5^*th*^ percentile of *X* and *y* is the 5^*th*^ percentile of *Y*

*RR*: *x* is the 5^*th*^ percentile of *X* and *y* is the 95^*th*^ percentile of *Y*

Here, the first annotation determines whether *x* is the 95^*th*^ (for *F*) or 5^*th*^ (for *R*) percentile of *X*, while the second annotation sets *y* ‘s percentiles reciprocally (*F*: 5^*th*^, *R*: 95^*th*^) regarding *Y*. The strand annotation depended percentile scheme is required due to the abovementioned mirroring of entry areas (in the context of the matrix folding scheme). Due to this mirroring, an entry can expand into an area in four different directions.

### Large insertions and the sentinel vertex

Let *I* be a large insertion and let *r* be a read that covers *I* partially at the beginning or end without fully enclosing *I*. In this case, an edge corresponding to *I* cannot be formed from *r* because either the edge’s origin vertex or destination vertex are unknown. Accordingly, it is impossible to create a valid adjacency matrix entry for *I* using *r*. For coping with this situation, a sentinel vertex and its mate (one for forward strand and one for reverse strand) are added to our graph model. These sentinels take the position of the missing origin or destination vertex in the context of the edge creation. As for all vertices, one row and one column of the folded adjacency matrix correspond to the sentinel vertex and its mate (see Supplementary Note 8).

Practically, edges are connected to the sentinel vertex or its mate if the outer ends of a read exceed a given distance to the outmost ends of its seeds (MEMs). Further, the previously introduced clustering for adjacency matrix entries can be used with the sentinels’ column and row as well. If there are reads that fully enclose *I*, a merging with entries that only partially enclose *I* is possible. If such fully enclosing reads are absent, genome assembly techniques are required for the reconstruction of *I* and its matrix entry. Supplementary Note 8 comprises a detailed description of the creation, clustering and merging of edges that connect to the sentinel vertex or its mate.

### Reconstructing sequenced genomes from graphs

Highly accurate SV calling should allow a reconstruction of the sequenced genome on the foundation of the reference genome and the SV calls. In the following, we assume the existence of a sequenced genome *S* together with a set *X* of simulated long (PacBio) and short (Illumina) reads for *S*. Using our proposed approach, we compute an adjacency matrix *M*_*S*_ using *S* as one single error-free read and a matrix *M*_*X*_ using the set of reads *X*. The matrices *M*_*S*_ and *M*_*X*_ can differ slightly due to the impact of sequencing errors, the size of reads in *X* and the repetitiveness of the reference genome. The differences between *M*_*S*_ and *M*_*X*_ can manifest in three ways: 1) An entry of *M*_*S*_ is slightly misplaced in *M*_*X*_. 2) An entry of *M*_*S*_ is missing in *M*_*X*_ and 3) *M*_*X*_ comprises an additional entry that does not occur in *M*_*S*_. The matrices *M*_*S*_ and *M*_*X*_ define two skew-symmetric genome-mapping graphs *G*_*S*_ and *G*_*X*_, respectively. Let *t*_*S*_ be a traversal through *G*_*S*_ that visits the edges in compliance with *S. t*_*S*_ defines a set of tuples *T*_*S*_ = {(*i, e*): *e* is the *i*th edge visited during the traversal *t*_*S*_}. Here an edge can be part of multiple tuples in *T*_*S*_. By following the edges in *T*_*S*_ in ascending order of their *i*-values, we can reconstruct *S* in terms of *G*_*S*_. In this context, the diagonal entries (which are not part of the matrix in practice for efficiency reasons) are implicitly inserted. According to the computation of *t*_*S*_ on the foundation of *S*, it must be possible to compute a traversal *t*_*X*_ through *G*_*X*_. For this purpose, the partial traversal information comprised in reads needs to be accumulatively combined using techniques known from genome assemblers. Further, insertions that are not fully covered by any read require read-to-read alignments for reconstruction. These two problems are subject to further research. In this work, we instead infer *T*_*X*_ directly from *T*_*S*_ as follows: For each pair (*i, e*) ∈ *T*_*S*_, we search for the spatially closest neighbor *e*′ ∈ *M*_*X*_. If the distance to *e*′ exceeds a given threshold, we ignore the pair (*i, e*). Otherwise, we add a tuple (*i, e*′) to an initially empty *T*_*X*_. In the context of this addition, we copy the weight (weights are insertions) of *e* to *e*’ too. Additionally, we define a successor function on the indices in *T*_*X*_ as *succ*(*i*) = min({*j* | *j* > *i* and (*j, e*) ∈ *T*_*X*_}). For example, in Supplementary Note 14, we have *succ*(1) = 2 and *succ*(2) = 4 for *T*_*X*_. The reconstruction of an approximated *S* via *T*_*X*_ follows the same algorithmic approach as described for *S* and *T*_*S*_. However, due to the differences between *M*_*S*_ and *M*_*X*_, we get an approximated form of *S* merely. As shown in Supplementary Note 14, these differences can cause a situation, where two tuples (*i, e*) and (*succ*(*i*), *e*′′) in *T*_*X*_ contradict the graph *G*_*X*_. (I.e., the origin of the edge *e*′′ cannot be reached from the destination of the edge *e*.) In this case, we continue the reconstruction at the origin of *e*′′ as soon as we reach the destination of *e*. For example, in Supplementary Note 14, this contradiction occurs between the edges *a*′ and *b*′ as well as *b*′ and *d*′. Therefore, the reconstruction comprises the segments before the destination of *a*′ and the segments following the origin of *d*′ merely. By choosing the origin of *d*′ instead of its destination, we avoid a bypassing of the insertion *I*.

As mentioned before, our skew-symmetric graph model unifies forward and reverse strand. The reconstruction process can encounter edges connecting vertices of opposite strands that we call crossing edges. Due to such crossing edges, the strand has to be tracked during reconstruction. In *T*_*S*_ this tracking is achieved by inverting a “current strand”-variable whenever the reconstruction passes crossing edges. (The current strand has to be considered during the abovementioned insertion of implicit edges; see Fig. 6 C, E.) However, in *T*_*X*_, the strand tracking can fail due to the disappearance of crossing edges. For tackling this shortcoming, we inspect *T*_*S*_ whenever we resolve one of the abovementioned contradictions between *T*_*X*_ and *G*_*X*_.

## Competing Interests

The authors declare that they have no competing interests.

## Code availability

An open-source prototype implementation of our approach van be found at https://github.com/ITBE-Lab/MA. The scripts used for performing all experiments are available at https://github.com/ITBE-Lab/MSV-EVAL.

## Notes

https://github.com/ITBE-Lab/MSV-EVAL

### Competing Interest Statement

The authors have declared no competing interest.

### Summary of Updates

Figures and subtitles updated. Minor typos fixed.

## References

1. Sedlazeck FJ, Rescheneder P, Smolka M, Fang H, Nattestad M, von Haeseler A, Schatz MC: Accurate detection of complex structural variations using single-molecule sequencing. Nature Methods 2018, 15(6):461–468.

2. Layer RM, Chiang C, Quinlan AR, Hall IM: LUMPY: a probabilistic framework for structural variant discovery. Genome biology 2014, 15(6):R84.

3. Chen X, Schulz-Trieglaff O, Shaw R, Barnes B, Schlesinger F, Källberg M, Cox AJ, Kruglyak S, Saunders CT: Manta: rapid detection of structural variants and indels for germline and cancer sequencing applications. Bioinformatics 2015, 32(8):1220–1222.

4. Rausch T, Zichner T, Schlattl A, Stütz AM, Benes V, Korbel JO: DELLY: structural variant discovery by integrated paired-end and split-read analysis. Bioinformatics 2012, 28(18):i333–i339.

5. Chong Z, Ruan J, Gao M, Zhou W, Chen T, Fan X, Ding L, Lee AY, Boutros P, Chen J et al: novoBreak: local assembly for breakpoint detection in cancer genomes. Nature Methods 2016, 14:65.

6. Nattestad M, Schatz MC: Assemblytics: a web analytics tool for the detection of variants from an assembly. Bioinformatics 2016, 32(19):3021–3023.

7. Fan X, Chaisson M, Nakhleh L, Chen K: HySA: a Hybrid Structural variant Assembly approach using next-generation and single-molecule sequencing technologies. Genome research 2017, 27(5):793–800.

8. Abo RP, Ducar M, Garcia EP, Thorner AR, Rojas-Rudilla V, Lin L, Sholl LM, Hahn WC, Meyerson M, Lindeman NI et al: BreaKmer: detection of structural variation in targeted massively parallel sequencing data using kmers. Nucleic acids research 2015, 43(3):e19–e19.

9. Choi M-H, Sohn J-i, Yi D, Menon AV, Kim YJ, Kyung S, Shin S-H, Na B, Joung J-G, Yoon S et al: Ultra- fast Prediction of Somatic Structural Variations by Reduced Read Mapping via Pan-Genome <em>k</em>-mer Sets. bioRxiv 2020:2020.2010.2025.354456.

10. Fang L, Hu J, Wang D, Wang K: NextSV: a meta-caller for structural variants from low-coverage long-read sequencing data. BMC bioinformatics 2018, 19(1):180.

11. Kuzniar A, Maassen J, Verhoeven S, Santuari L, Shneider C, Kloosterman WP, de Ridder J: sv-callers: a highly portable parallel workflow for structural variant detection in whole-genome sequence data. PeerJ 2020, 8:e8214.

12. Cameron DL, Di Stefano L, Papenfuss AT: Comprehensive evaluation and characterisation of short read general-purpose structural variant calling software. Nature communications 2019, 10(1):1–11.

13. Kosugi S, Momozawa Y, Liu X, Terao C, Kubo M, Kamatani Y: Comprehensive evaluation of structural variation detection algorithms for whole genome sequencing. Genome Biology 2019, 20(1):117.

14. Mahmoud M, Gobet N, Cruz-Dávalos DI, Mounier N, Dessimoz C, Sedlazeck FJ: Structural variant calling: the long and the short of it. Genome Biology 2019, 20(1):246.

15. Chander V, Gibbs RA, Sedlazeck FJ: Evaluation of computational genotyping of structural variation for clinical diagnoses. GigaScience 2019, 8(9).

16. Pevzner P, Tesler G: Genome rearrangements in mammalian evolution: lessons from human and mouse genomes. Genome Res 2003, 13(1):37–45.

17. Hickey G, Paten B, Earl D, Zerbino D, Haussler D: HAL: a hierarchical format for storing and analyzing multiple genome alignments. Bioinformatics 2013, 29(10):1341–1342.

18. Needleman SB, Wunsch CD: A general method applicable to the search for similarities in the amino acid sequence of two proteins. Journal of molecular biology 1970, 48(3):443–453.

19. Smith TF, Waterman MS: Identification of common molecular subsequences. Journal of molecular biology 1981, 147(1):195–197.

20. Li H: Minimap2: pairwise alignment for nucleotide sequences. Bioinformatics 2018, 1:7.

21. Schmidt M, Heese K, Kutzner A: Accurate high throughput alignment via line sweep-based seed processing. Nature Communications 2019, 10(1):1939.

22. Yue J-X, Li J, Aigrain L, Hallin J, Persson K, Oliver K, Bergström A, Coupland P, Warringer J, Lagomarsino MC et al: Contrasting evolutionary genome dynamics between domesticated and wild yeasts. Nature genetics 2017, 49(6):913–924.

23. Zook JM, Hansen NF, Olson ND, Chapman L, Mullikin JC, Xiao C, Sherry S, Koren S, Phillippy AM, Boutros PC: A robust benchmark for detection of germline large deletions and insertions. Nature biotechnology 2020:1-9.

24. Collins RL, Brand H, Karczewski KJ, Zhao X, Alföldi J, Francioli LC, Khera AV, Lowther C, Gauthier LD, Wang H et al: A structural variation reference for medical and population genetics. Nature 2020, 581(7809):444–451.

25. Werling DM, Brand H, An J-Y, Stone MR, Zhu L, Glessner JT, Collins RL, Dong S, Layer RM, Markenscoff-Papadimitriou E et al: An analytical framework for whole-genome sequence association studies and its implications for autism spectrum disorder. Nature Genetics 2018, 50(5):727–736.

26. Collins RL, Brand H, Redin CE, Hanscom C, Antolik C, Stone MR, Glessner JT, Mason T, Pregno G, Dorrani N et al: Defining the diverse spectrum of inversions, complex structural variation, and chromothripsis in the morbid human genome. Genome Biology 2017, 18(1):36.

27. Danecek P, Auton A, Abecasis G, Albers CA, Banks E, DePristo MA, Handsaker RE, Lunter G, Marth GT, Sherry ST et al: The variant call format and VCFtools. Bioinformatics (Oxford, England) 2011, 27(15):2156–2158.

28. Nattestad M, Goodwin S, Ng K, Baslan T, Sedlazeck FJ, Rescheneder P, Garvin T, Fang H, Gurtowski J, Hutton E: Complex rearrangements and oncogene amplifications revealed by long-read DNA and RNA sequencing of a breast cancer cell line. Genome research 2018, 28(8):1126–1135.

29. Gotoh O: Optimal sequence alignment allowing for long gaps. Bulletin of mathematical biology 1990, 52(3):359–373.

30. Ohlebusch E, Abouelhoda MI: Chaining algorithms and applications in comparative genomics. Handbook of Computational Molecular Biology 2006.

31. Li H: Aligning sequence reads, clone sequences and assembly contigs with BWA-MEM. arXiv preprint 13033997 2013.

32. Kutzner A, Kim P-S, Schmidt M: A performant bridge between fixed-size and variable-size seeding. BMC Bioinformatics 2020, 21(1):328.

33. Jeffares DC, Jolly C, Hoti M, Speed D, Shaw L, Rallis C, Balloux F, Dessimoz C, Bähler J, Sedlazeck FJ: Transient structural variations have strong effects on quantitative traits and reproductive isolation in fission yeast. Nature Communications 2017, 8(1):14061.

34. Homer N: Dwgsim: whole genome simulator for next-generation sequencing. GitHub repository 2010.

35. Zook JM, Catoe D, McDaniel J, Vang L, Spies N, Sidow A, Weng Z, Liu Y, Mason CE, Alexander N et al: Extensive sequencing of seven human genomes to characterize benchmark reference materials. Scientific Data 2016, 3:160025.

36. Goldberg AV, Karzanov AV: Path problems in skew-symmetric graphs. Combinatorica 1996, 16(3):353–382.

37. Roberts M, Hayes W, Hunt BR, Mount SM, Yorke JA: Reducing storage requirements for biological sequence comparison. Bioinformatics 2004, 20(18):3363–3369.

