## Supplementary Notes for "State-of-the-art structural variant calling: What went conceptually wrong and how to fix it?"

#### Supplementary Note 1. Breakends versus Breakpoints

Often the notion ‘breakpoint’ is used in the context of SV calling for identifying positions on the reference genome and reads [6, 7]. The number of breakpoints varies for different types of SV. For example, an insertion has two breakpoints on the read and one breakpoint on the reference, while a deletion has one breakpoint on the read and two breakpoints on the reference. In the context of our proposed approach, we aim for a description of SV that does not use atomic SVs. Therefore, we need terminology that expresses the endpoints of SV independent of their type. For this purpose, we rely on the notion ‘breakend’ as introduced in the “Specifying complex rearrangements with breakends” section of the “The Variant Call Format (VCF) Version 4.2 Specification”. In contrast to breakpoints, two breakends can occur in the same position on reference or read. Accordingly, all atomic SVs are characterized by exactly four breakends (two on the read and two on the reference). Here we use the notion breakend pair for referring to the respective two breakends on read or reference.

#### Supplementary Note 2. Diagrammatic dot-plots and genome section pictograms

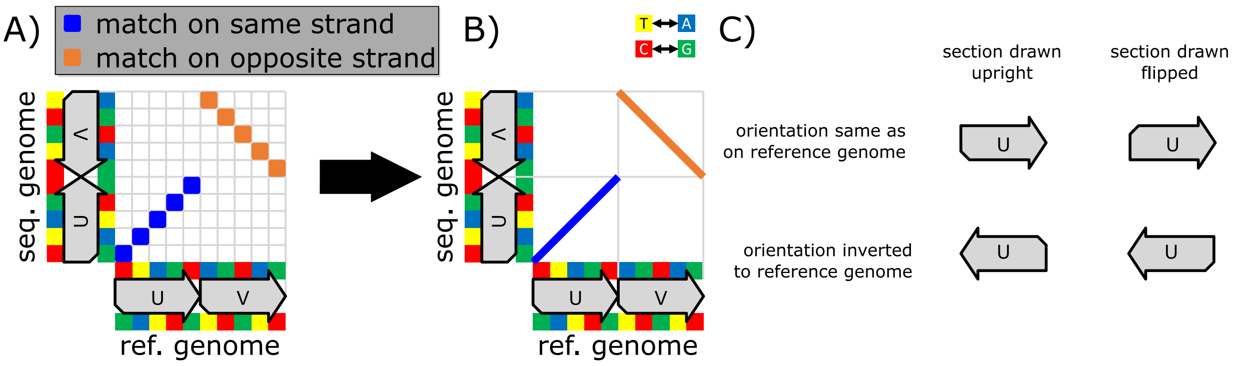

Subfigure **A)** shows a dot-plot, where each blue box indicates a match on identical strands and each orange box indicates a match on opposite strands. The reference genome and the sequenced genome appear on the x-axis and y-axis, respectively. For the reference genome, the colored squares on top of the sections $U, V$ represent the nucleotides on the forward strand. Accordingly, the reverse strand appears below the sections. The forward strand of the sequenced genome is on the left of its genome section, while the reverse strand is on its right.

Subfigure **B)** displays a diagrammatic representation of the left side's dot-plot. In this diagrammatic representation, consecutive matches (on equal as well as opposite strands) appear as lines, where the start of the line includes the bottommost match and the end includes the topmost match. The yellow, red, blue and green squares on the reference genome and sequenced genome represent the nucleotides T, C, A and G, respectively.

Subfigure **C)** explains our visualization for genome sections using arrow-like pictograms. A pictogram can comprise a nook, which indicates that its nucleotide sequence occurs on the reference. This nook in combination with the arrowheads expresses the direction of the pictogram's nucleotide sequence on the reference genome. Via this scheme, we visualize that genomic inversions are $180°$ rotations (and not a mirroring operation).

#### Supplementary Note 3. Unfolded matrices and full graphs for Fig. 1 of the main text

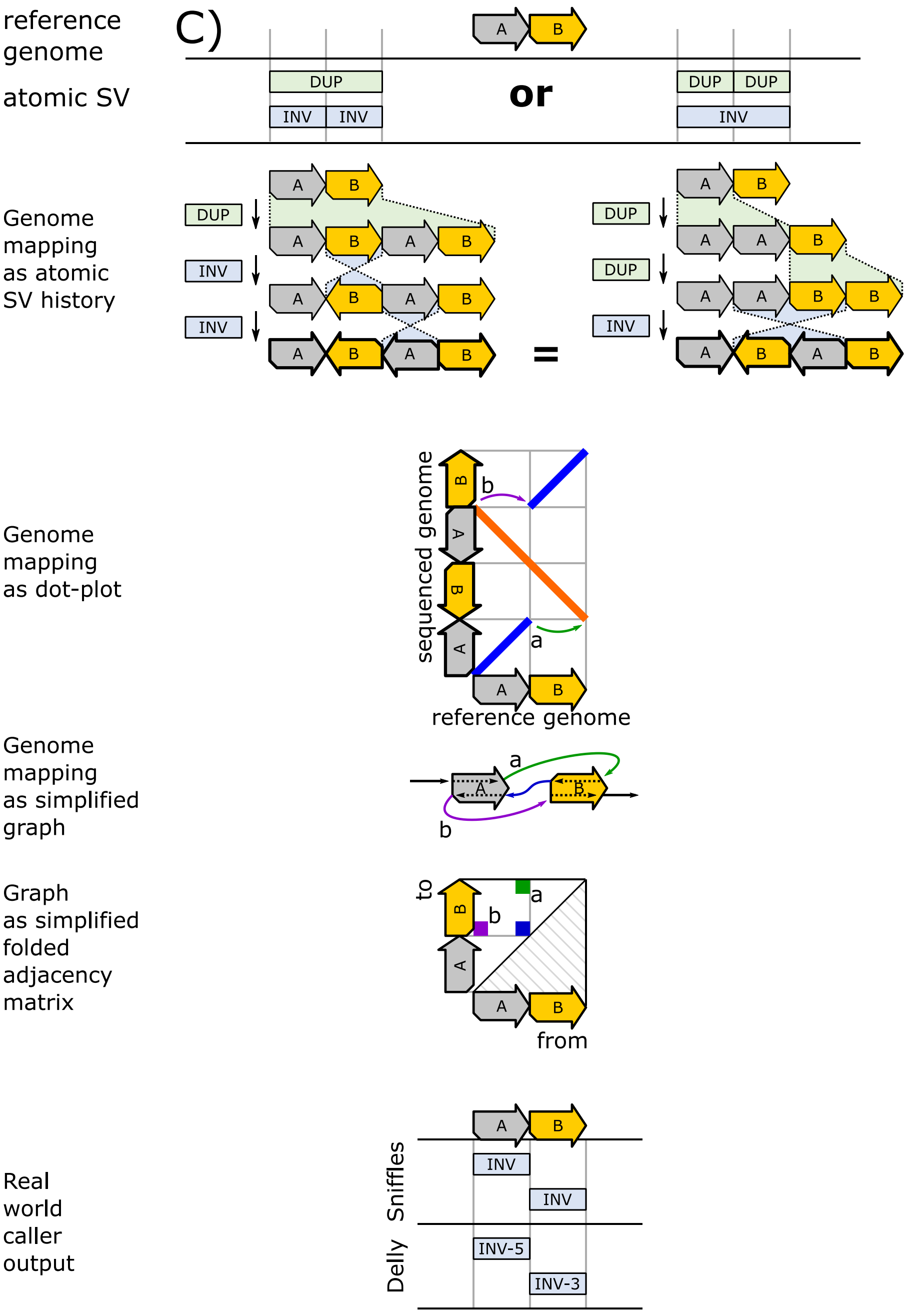

Supplementary Figure 3.1: The figure shows that a duplication followed by two inversions can lead to the same genomic outcome as two duplications followed by one inversion. The respective unfolded adjacency matrix and the full graph are displayed in the figure below.

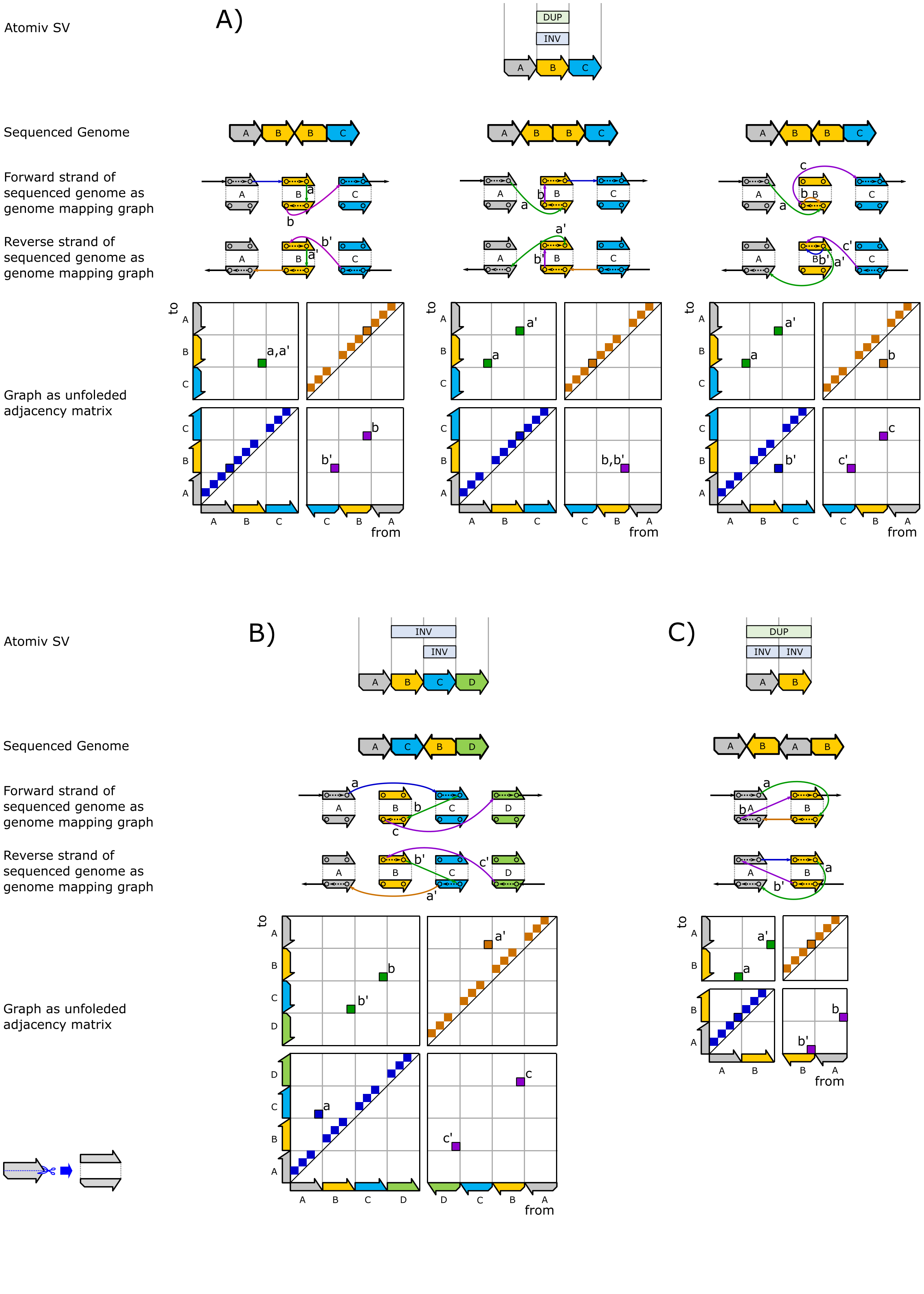

Atomiv SV
A)
DUP
INV
Sequenced Genome
Forward strand of sequenced genome as genome mapping graph
Reverse strand of sequenced genome as genome mapping graph
Graph as unfoleded adjacency matrix
to
from
B)
INV
INV
C)
DUP
INV
INV

Supplementary Figure 3.2: Fig. 1 of the main text depicts several ambiguities inherent to the description of genomic rearrangements using atomic SV. The main text's figure merely shows simplified instances of our graph model and their folded adjacency matrices.

The above figure displays the unfolded matrices together with their respective full skew-symmetric graphs. Subfigure A) and B) correspond to Fig. 1 A) and B) of the main text, respectively. Subfigure C) corresponds to Supplementary Figure 3.1. The folding scheme for matrices is described in the methods section of the main text. Black outlined entries correspond to the equally colored edges of their respective graph. All other entries belong to dashed edges (within the genome sections $A, B, C, D$) and are not shown in Fig. 1 of the main text.

Due to the folding scheme for matrices, the outlined blue and orange matrix entries in C) correspond to the single unlabeled blue entry in the respective folded matrix. There, the “from-to” direction of this unlabeled blue entry does not match the direction of the corresponding edge in the simplified graph. The same apparent paradox affects the edge labeled $b$ in Fig. 1 B). The unfolding of the matrix always resolves such paradoxes (see the above subfigure B and C).

#### Supplementary Note 4. Experimental setup for the generation of nested genomic rearrangements

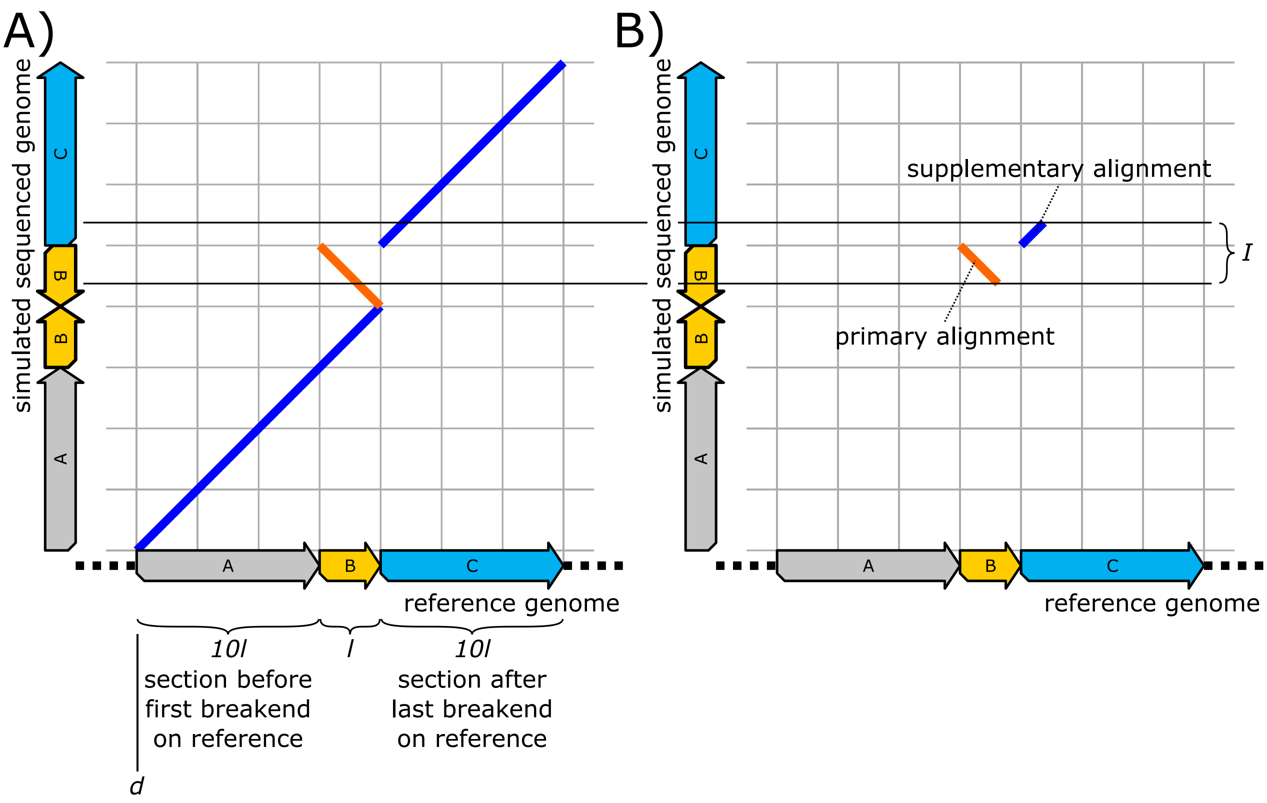

Ambiguities that are inherent to the description of genomic rearrangements via atomic SV are evaluated by applying the following four steps:

(1) A specific (generated) genomic rearrangement is represented using a diagrammatic dot-plot.

(2) The resulting diagrammatic dot-plot is turned into error-free alignments (incl. CIGAR).

(3) The error-free alignments are forwarded to various SV callers (Sniffles etc.) for SV detection.

(4) The detected SVs of the respective SV callers are evaluated and visualized.

These steps are explained in detail now:

1) We use the human genome (GRCh38.p12) as the reference genome; the length parameter $l$ (see above figure) is set to $1000$ nt. The length of the sections labeled $A, B, C$ of Fig. 1 of the main text are equal to the sizes of the respective sections in the above figure (although differently visualized in Fig. 1). The increased size of the sections $A$ and $C$ compared to $B$ creates SV-free regions for supporting the sampling step that is part of some SV callers (e.g. Delly). Each diagrammatic dot-plot is manually translated into a set of pseudo-seeds $X$ that incorporates the dot-plot's genomic rearrangement, where a pseudo-seed is

a quadruple $(q, r, l, \xi)$ of a query position $q$, reference position $r$, length $l$ and a value $\xi \in \{same, opposite\}$ that indicates equivalence between same or opposite strands. (Same strand and opposite strand pseudo-seeds are visualized as blue lines and orange lines, respectively.) All manually created pseudo-seeds are maximally extended in either direction. After creation, all pseudo-seeds in $X$ are moved by a random distance $d$ on the reference genome's first chromosome. For the above example, this delivers the set $Y = \{(0, d, 11l, same), (11l, 10l + d, l, opposite), (12l, 11l + d, 10l, same)\}$. The resulting set $Y$ defines the simulated sequenced genome. Insertions are injected by creating random sequences (Mersenne Twister random generator) of appropriate size. In the above example, the simulated sequenced genome is $AB\tilde{B}C$ for the reference genome $ABC$.

2) For generating alignments, intervals are picked on the simulated sequenced genome from 1). For long-read alignments, these intervals are of size $2 * l - 1$ (so that alignments cover no more than two breakend-pairs). For paired short read alignments, we rely on interval pairs, where each interval has a size of 250 nt and the distance between a pair is 100 nt. Using a sequenced genome interval $I$, an alignment is computed as follows:

We create a copy $Y'$ of $Y$ and limit the pseudo-seeds in $Y$' to the interval $I$ afterward. For this purpose, we delete all pseudo-seeds in $Y'$ that are completely outside of $I$ and shorten seeds that cross $I$'s endpoints. The remaining pseudo-seeds in $Y'$ are used for the creation of alignments. Here each seed results in a single alignment with a CIGAR comprising a match equally sized to the length of the pseudo-seed. Among these, all alignments except for the longest one are marked as supplementary. The reference position of the alignments is given by their pseudo-seed's reference positions. The query (read) position of the alignments is the relative distance of their pseudo-seeds to the beginning of $I$. The simulated read is the sequence in $I$ on the simulated sequenced genome. The strand of an alignment is given by the strand of its pseudo-seed. The above scheme for alignments and reads generation is used for generating a coverage of 100 (within the reference genome's section $ABC$).

3) All alignments generated in step 2) are collected in a SAM file that is forwarded to the respective SV caller.

4) By reading the fields 'ALT', 'POS' and 'INFO->END', the VCF-output of the SV caller is visualized as labeled intervals on the reference genome. Here the order in the diagrams is equal to the order of their respective occurrence in the VCF-file.

Our prototype (see Supplementary Note 6) computes all folded matrices exactly as shown in Fig. 1 of the main text.

#### Supplementary Note 5. SVs that are hidden to aligners – Extended analysis

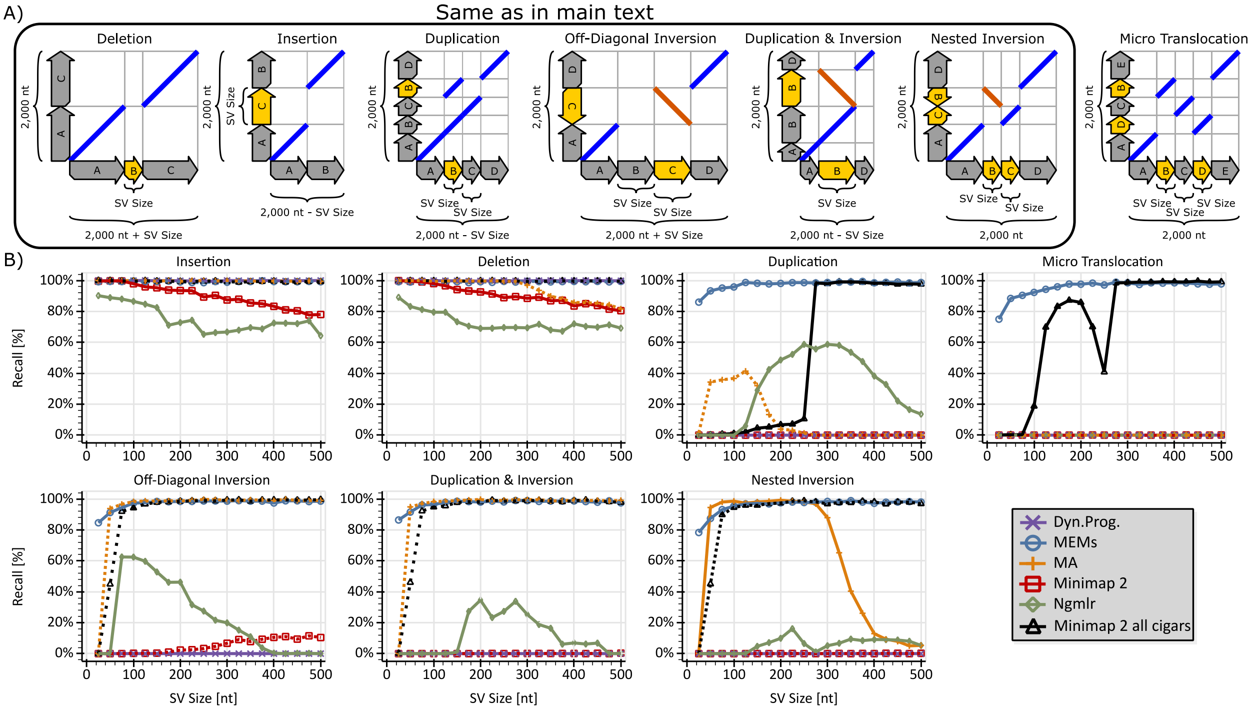

In addition to Fig. 2 of the main text, the above figure analyses micro translocations and comprises curves that display the recall rates for Minimap2 with a special setup that avoids discarding chains. The detailed parameters for Minimap2's setup are: "-z 400,1 --splice -P". The setting of the "-z" parameter to "400,1" decreases the z-drop, which, in turn, improves the discovery of inversions at the cost of increased runtime and more false positives. The "--splice" parameter enables splice mode. The "-P" instructs Minimap2 to keep all chains for the price of tremendously increased runtimes and excessively large SAM files.

For micro translocations, all aligners fail except Minimap2 with the above special setup. This behavior can be explained by the path-contradicting nature of this kind of genomic rearrangement. The different behavior of Minimap2 for the two settings (red curve and black curve) indicates that, for getting reasonable runtime, Minimap2's heuristics (z-drop, chaining, etc.) remove information vital to the discovery of genomic rearrangements.

#### Supplementary Note 6. Computation of ground truth entries

The computation of ground truth entries happens by applying the entry creation scheme described in the methods section with the sequenced genome as one single error-free read. It consists of the following three steps:

1. Obtain all MEMs using Minimizers and occurrence filtering as described in the main text.
   (see "Incorporating genome repetitiveness and sequencer errors into SV calling" in the results section.)
2. Apply an overlap elimination on the MEMs as described in the main text.
   (see "Incorporating genome repetitiveness and sequencer errors into SV calling" in the results section.)
3. Turn the MEMs into matrix entries as described in the main text.
   (see "SV calling using raw MEMs" in the results section.)

The matrix entries resulting from the final step represent the ground truth. Because the sequenced genome acts as one long perfect read, additional entry processing (e.g. fuzzy inference of edges, computing clusters by merging overlapping entry-areas and approximating true entry locations) is not required here.

For the genome reconstruction process (see Supplementary Note 14), we additionally memorize the visiting order of the matrix entries in a traversal (through the graph corresponding to the matrix) that delivers the sequenced genome. This visiting order is equivalent to the occurrence order of the corresponding MEMs on the sequenced genome.

#### Supplementary Note 7. Description of the SV caller prototype and benchmarking environment

All evaluation and analysis of our approach is performed by using a prototype implementation that is characterized as follows:

- Similar to the approach of the Modular Aligner MA [1] and TensorFlow [2], Python 3 is used for constructing a computational graph that redirects all computational operations to a C++ library (realized in C++-17). This architecture keeps Python inactive during graph execution for gaining high performance and exploiting parallel computing (multithreading) advantages.
- For gaining a high level of flexibility, the C++ library, in turn, relies on a PostgreSQL [3] database as a backend. The database is used for storage purposes (e.g. reads, adjacency matrix) and efficient data retrieval operations using SQL [4].
- For visualizing purposes, an additional viewer application is realized using the Python-Bokeh library [5]. On the database level, the viewer exploits the highly efficient storage and retrieval of spatial data offered by PostGIS.
- The prototype is publically available on GitHub under the MIT license on https://github.com/ITBE-Lab/MA
- Our evaluation framework is publically available on GitHub under the MIT license on https://github.com/ITBE-Lab/MSV-EVAL

All experimental work is done on an *AMD Ryzen Threadripper 1950X 16-Core Processor* with 128 GB RAM.

The versions of third-party software and applications are listed below:

- Minimap2 2.17 (r941)
- Delly v0.8.1
- Sniffles 1.0.8
- NGMLR 0.2.8
- Postgres 12.3
- Postgis 3
- MA 2.0.0-afc9ac7

#### Supplementary Note 8. The sentinel vertex

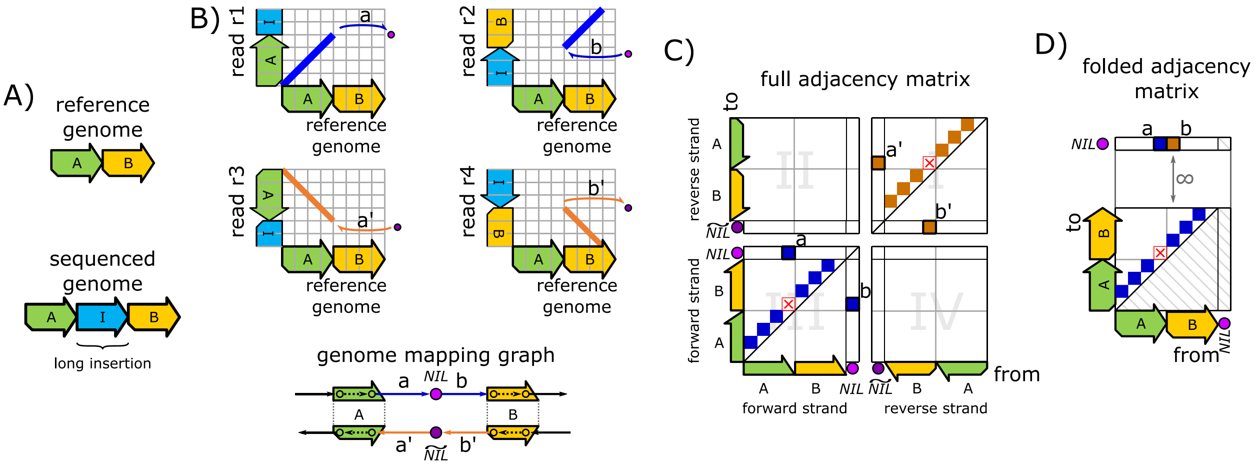

Supplementary Figure 8.1: **A)** displays a sequenced genome $AIB$ and a reference genome $AB$. $I$ is assumed to be a long insertion that is not fully enclosed by any read. **B)** shows four reads that partially cover $I$ and their genome mapping graph. The two reads $r1$, $r2$ originate from the forward strand, while the remaining two reads $r3$, $r4$ originate from the reverse strand of the sequenced genome. The reads $r1$, $r3$ cover the beginning of $I$, while the reads $r2$, $r4$ cover the end of $I$. All four reads create an edge that connects to the sentinel vertex or its mate (labeled $NIL$ and $\widetilde{NIL}$ in the genome mapping graph). **C)** presents the unfolded adjacency matrix for the genome mapping graph in B). Matrix entries without black outlines are implicit (see methods section of main text). Red crosses mark the positions of entries for reads that would fully enclose the insertion. These crosses are in line (either horizontally or vertically) with the entries that connect to sentinel vertices (because their origin or destination vertices are equal) and they can be used for merging these entries. **D)** Forward strand and reverse strand are unified by applying our folding scheme to the adjacency matrix (the methods section of the main text describes this folding scheme). Because of this folding, there are no entries in the $NIL$ column on the right. For applying our clustering to sentinel connecting entries, the sentinel's row is placed far above the remaining matrix. This prevents clusters of sentinel entries from interfering with clusters of regular entries.

In the following, our approach is extended by the concept of the sentinel vertex for coping with long insertions:

##### Creation of sentinel entries in the adjacency matrix

The adjacency matrix is extended by a column and row labeled $NIL$ as well as a column and row labeled $\widetilde{NIL}$ for the sentinel vertex and its mate. These inserted rows and columns are placed in central locations of the unfolded adjacency matrix as shown in Supplementary Fig. 8.1. In the folded matrix, the sentinels correspond to the topmost row and rightmost column.

Let $(q, r, l, k)$ be a seed with the beginnings $q$ and $r$ on a query $Q$ (read) and reference (genome) as well as a length $l$ and a strand information $k \in \{F, R\}$, where $F$ and $R$ correspond to forward and reverse strand, respectively. Further, let $d$ be a given minimal distance (threshold) that triggers the connection to a sentinel. Sentinel related matrix entries are created as follows:

| conditions | | | outcome |
|---|---|---|---|
| **strand information ($\boldsymbol{k}$)** | **first or last seed on read** | **distance to end of read** | **entry in adjacency matrix** |
| $F$ | first (smallest $q$) | $q \geq d$ | $(NIL, r)$ |
| $R$ | first (smallest $q$) | $q \geq d$ | $(\widetilde{NIL}, r + l)$ |
| $F$ | last (largest $q + l$) | $q + l \leq \lvert Q \rvert - d$ | $(r + l, NIL)$ |
| $R$ | last (largest $q + l$) | $q + l \leq \lvert Q \rvert - d$ | $(r, \widetilde{NIL})$ |

By applying the matrix-folding scheme described in the main text, sentinel entries from forward and reverse strand reads are unified. The distance $d$ must be chosen according to the specific characteristics of a set of reads as e.g. the error-rate of the sequencer and average length of the reads. In the context of our evaluation of yeast genomes, we use a distance $d = 50$ for CCS PacBio reads and $d = \infty$ for Illumina reads, where $\infty$ disables the creation of sentinel connecting entries.

#### Clustering of sentinel connecting entries

The matrix entries' clustering described in the methods sections exploits the spatial locality of the vertices (corresponding to the entries) on the reference genome. This spatial locality is void between the sentinels and all other vertices. Therefore, clusters of sentinel connecting entries are not allowed to overlap with clusters of regular entries. As shown in Supplementary Fig. 8.1 D), we separate both types of clusters by inserting a void space between them in the adjacency matrix.

#### Merging of sentinel connecting entries with regular adjacency matrix entries

In the following, Let $X$ be the set of all reads that cover some section of an insertion $I$. We split the set $X$ into three subsets $X_{Full}$, $X_{In}$, $X_{Out}$ as follows: $X_{Full}$ comprises all reads in $X$ that fully enclose the insertion $I$. $X_{In}$ consists of all reads in $X$ that cover the beginning of $I$ but do not fully enclose $I$. Accordingly, $X_{Out}$ keeps all reads of $X$ that cover the end of $I$ but do not fully enclose $I$.

For all reads in $X_{Full}$, the resulting entries for $I$ are regular entries that are created via the breakends of two seeds. In contrast, the reads in $X_{In}$ and $X_{Out}$ connect to the sentinels for $I$. However, the origin of the entries in $X_{In}$ matches the origin of the entries in $X_{Full}$ and the destination of the entries in $X_{Out}$ matches the destination of the entries in $X_{Full}$. Via these relationships, it is possible to merge the entries of all three subsets to one single matrix entry. In the context of a matrix entry's scoring, this merging is exploited for measuring its number of supporting reads.

### Supplementary Note 9. Ambiguities inherent to the state-of-the-art SV calling

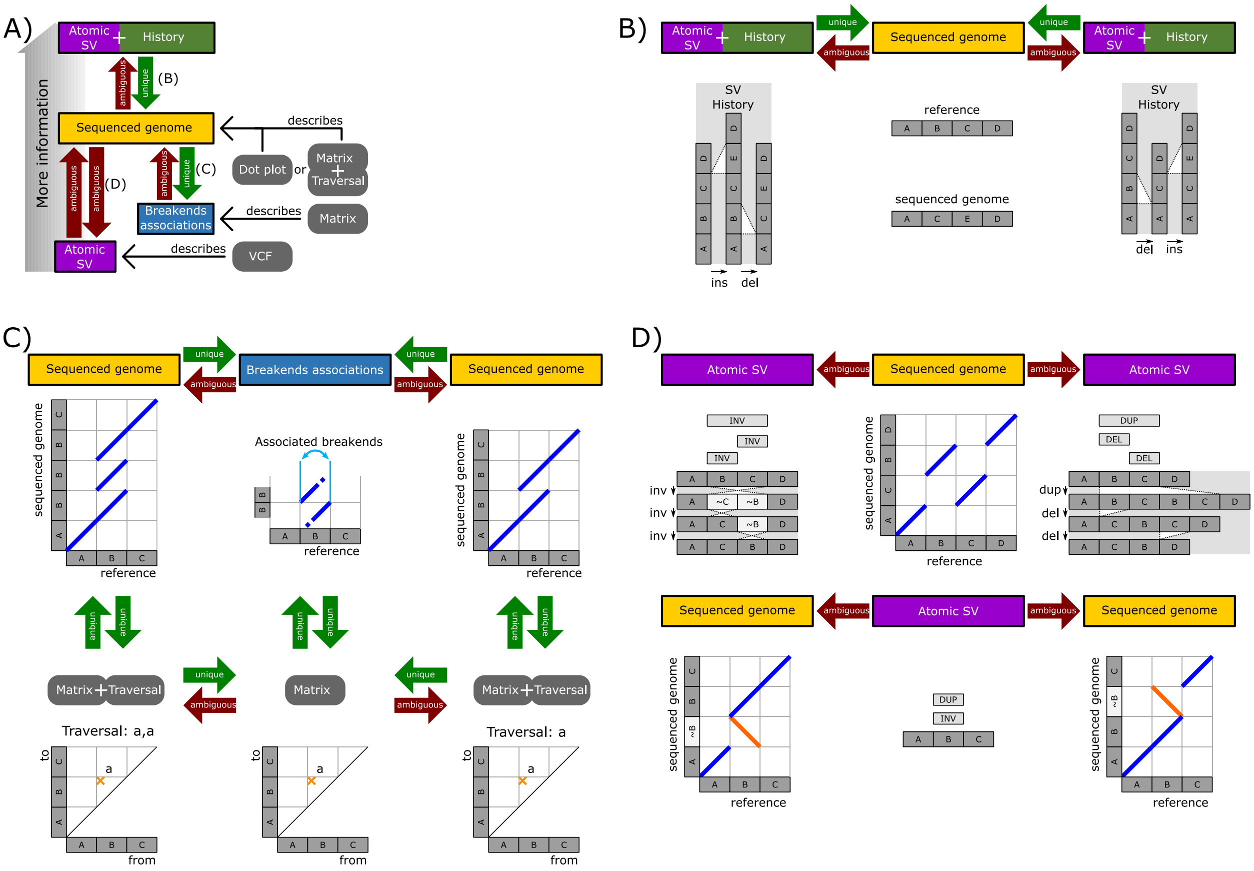

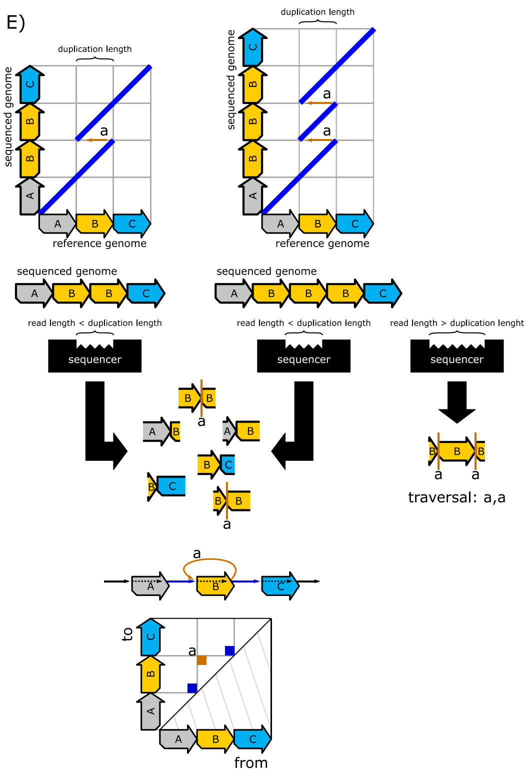

The above figure shows examples for the ambiguities mentioned in the discussion section and Fig. 5 of the main text. **A)** is a copy of Fig. 5 with additional annotations that indicate the locations of the cases B) to D) regarding the remainder of the above figure.

Subfigure **B)** shows that even in simple cases with isolated atomic SV, the history (order of occurrence) cannot be reconstructed from the sequenced genome.

**C)** The left and right sequenced genome have identical associated breakends (center column) on the reference genome. However, the left sequenced genome has two duplications of $B$ while the right sequenced genome comprises one duplication merely. Our approach catches the equivalence of the associated breakends via a corresponding equivalence in the adjacency matrix, while the difference in the number of occurrence of $B$ corresponds to two different graph traversals. The separation of adjacency matrix (breakend associations) and traversal (order of breakend associations on the sequenced genome) is mandatory due to the limited size of real-world reads (see subfigure E)).

**D)** In the top row, the sequenced genome $ACBD$ is created using two different sets of atomic SV from the reference genome $ABCD$. Accordingly, in the bottom row, a duplication and an inversion on the section $B$ are used for creating two different sequenced genomes. Notably, the atomic SVs are applied in the same order for both outcomes.

**E)** Each read that does not span the full genome delivers a partial graph traversal merely. Often, these partial graph traversals cannot be combined into a full traversal. Particularly with short reads, such a combination is often impossible in real-world scenarios.

#### Supplementary Note 10. Detailed matrix folding of Fig. 6 in the methods section

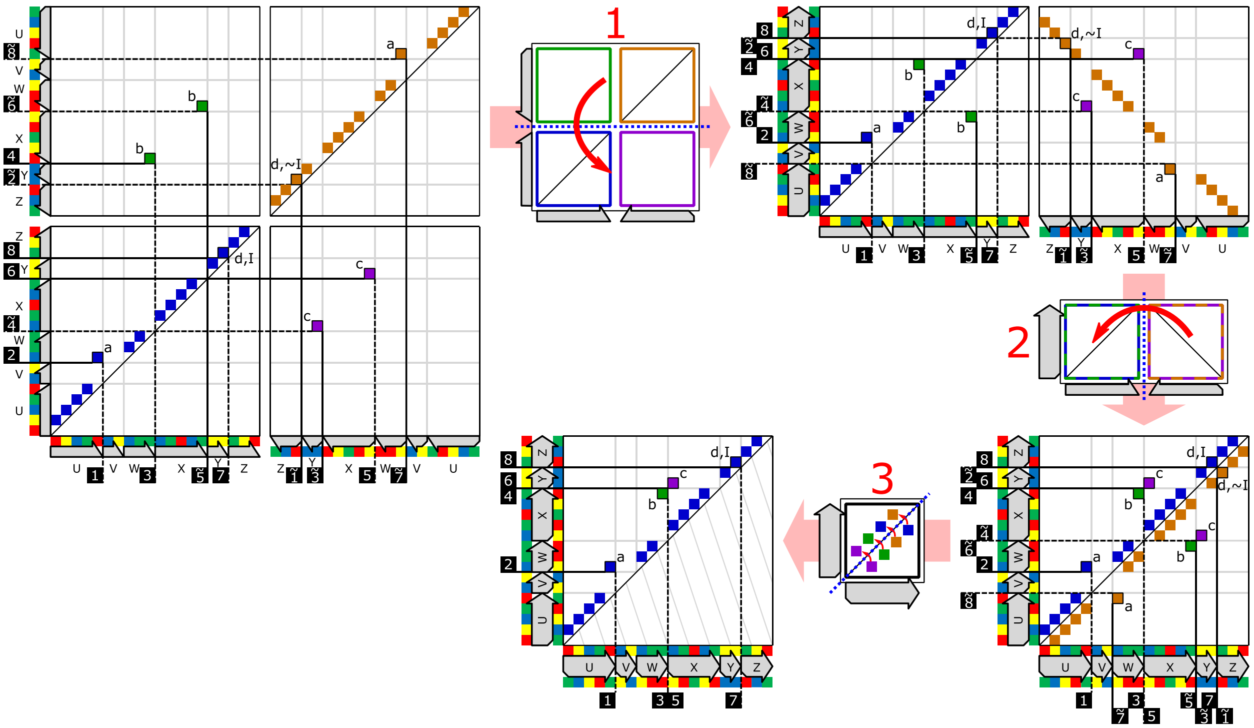

The above figure shows the detailed folding of the adjacency matrix in Fig. 6 E) in the main text.

#### Supplementary Note 11. Detailed description of overlap-elimination

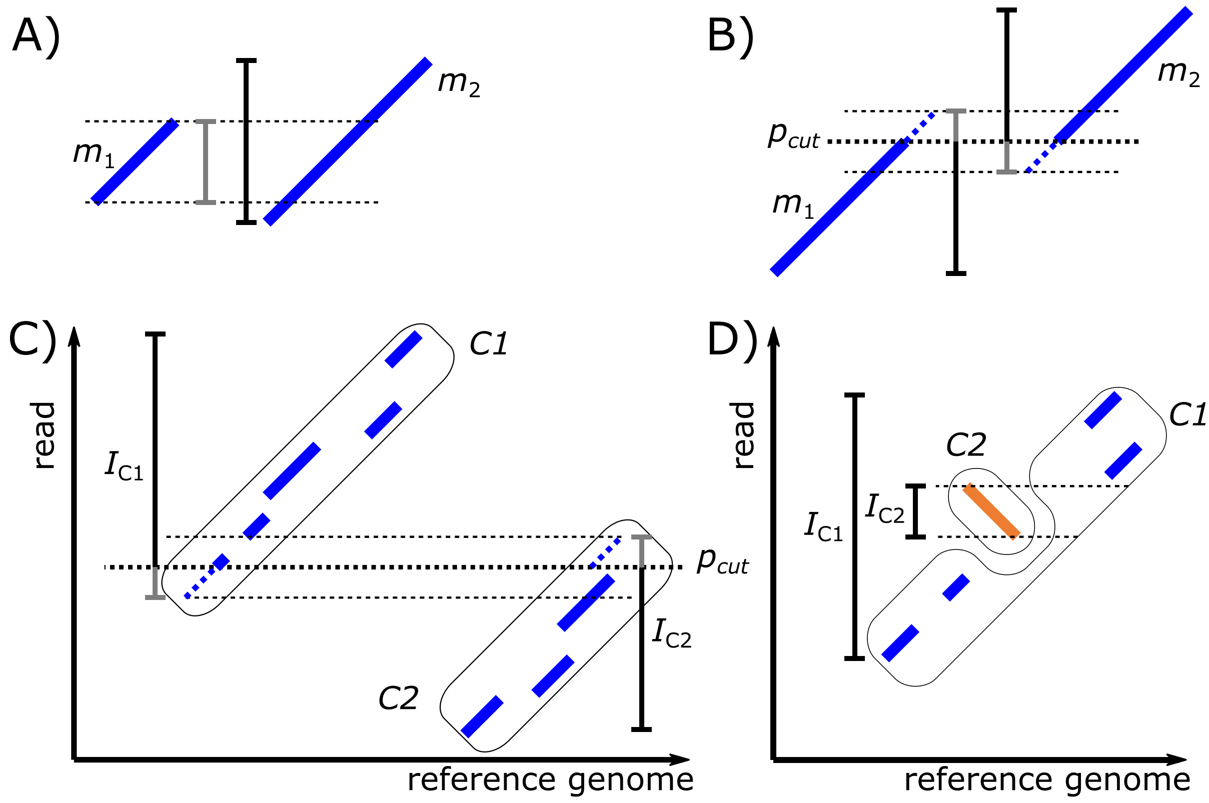

Figure 3 D) of the results section displays a situation, where two MEMs overlap on the y-axis of a diagrammatic dot-plot. “Computing MEMs for a read via reseeding” of the methods section introduces an overlap elimination technique for managing such overlaps. Subfigure **A)** and **B)** illustrate the cutting scheme described there. The labels $m_1$, $m_2$ and $p_{cut}$ correspond to the two MEMs and to the central cutting location mentioned in the main text, respectively.

If the outer ends of large-scale SV belong to repetitive regions on the reference genome, the overlaps visualized in **C)** between two clusters of MEMs occur. Here, an application of the overlap elimination on MEMs removes all overlaps but connects the repetitive regions with several entries in both directions. However, the repetitive regions should be connected by one entry merely. For solving this problem, overlap elimination is applied to MEM-clusters as well. For MEM-cluster creation, we rely on Strips of Consideration [1]. Let $C$ denote a cluster and $(q, r, l)$ a MEM in $C$, where $q, r$ and $l$ are the start position on query, the start position on reference and length of the MEM, respectively. The read interval of $C$ is defined as

$$I_C := [\min\{q | (q, r, l) \in C\}, \max\{q + l | (q, r, l) \in C\}).$$

The overlap elimination scheme for MEMs as described in the main text is applied to MEM-clusters using their respective read intervals. **D)** visualizes one special case for enclosed intervals. If the enclosing MEM-cluster and the enclosed cluster do not comprise overlapping seeds, both clusters are kept.

#### Supplementary Note 12. Fuzzy inference of edges from MEMs

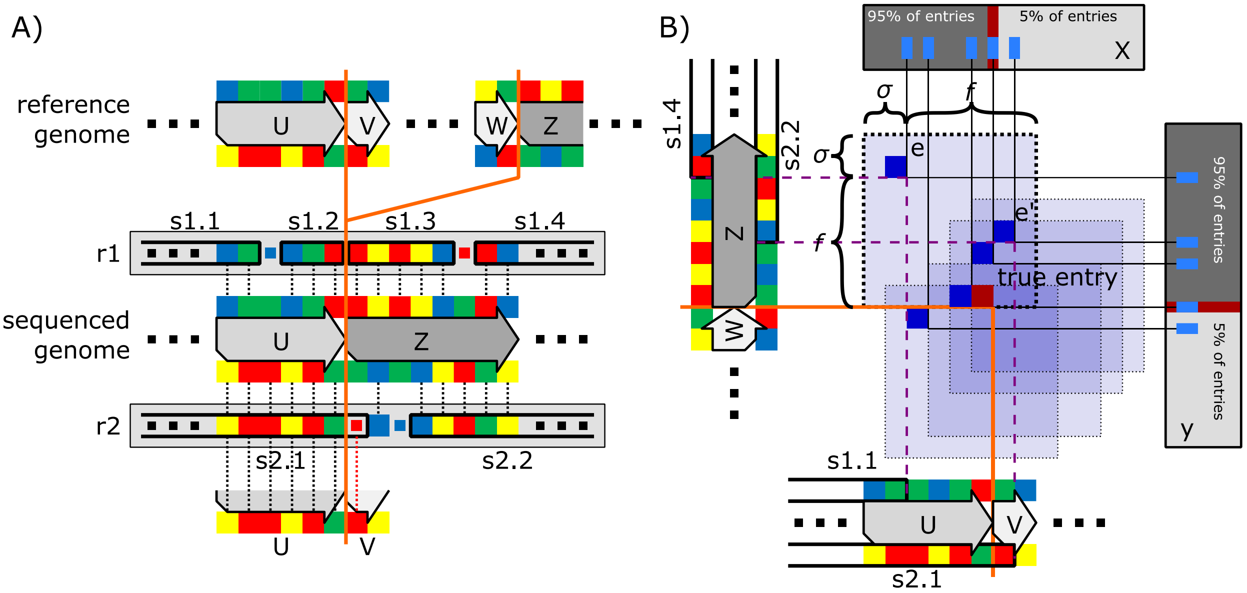

The figure explains the impact of sequencing errors on matrix entries. **A)** In $r1$, two sequencing errors cause the MEMs $s1.1$ and $s1.4$ to break prematurely regarding the breakend pair that is visualized as an orange line. As a result, the two short MEMs $s1.2$ and $s1.3$ appear. We assume that these two MEMs get lost during the occurrence filtering because of their small size. Due to the absence of these two MEMs, we get the entry $e$ in B). In $r2$, the first sequencing error causes $s2.1$ to erroneously extend over the breakend on the sequenced genome, while the second sequencing error causes a premature breaking of $s2.2$. The two MEMs $s2.1$ and $s2.2$ create the entry $e'$ in B). **B)** shows the entry areas in the folded adjacency matrix for the entries $e$ and $e'$ as well as three more fictional entries. Entry areas are visualized as light blue squares. Further, the subfigure indicates the location of the true entry via a red box (matrix entry). The distances $\sigma$ and $f$ are expressed with respect to the entry $e$. The gray shaded sidebars labeled $X$ and $Y$ display the two sets used for the approximation of the true entry's location. In this context, the red lines in the sidebars visualize the true entry location according to our proposed percentile scheme (5% to 95%).

#### Supplementary Note 13. Line-sweep based clustering of entry-areas

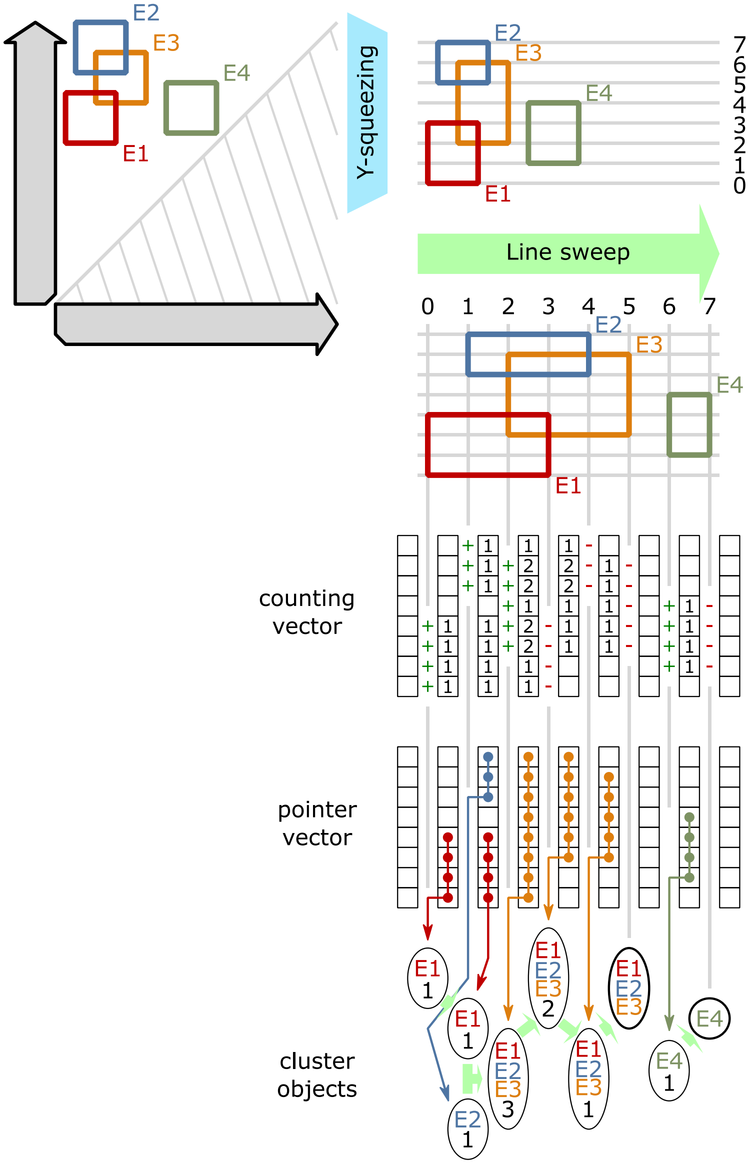

Practically, we store entry-areas as a list of rectangles, where the same rectangle can occur multiple times. By sorting this list followed by a line-sweep, we can identify overlapping entry-areas in the matrix. This line-sweep based clustering starts with an inexact sweep that separates the whole dataset into smaller inexact clusters. Here, an inexact cluster may be comprised of several actual clusters (i.e. sets of rectangles that actually overlap), but actual clusters always belong to a single inexact cluster. Given an inexact cluster, we subsequentially compute the actual clusters within it using an exact line-sweep. Theoretically, the exact line-sweep is capable of computing the exact clusters directly from the complete dataset; however, this would increase memory requirements and runtime tremendously.

#### The exact algorithm

We start by explaining the exact line-sweep: The algorithm sweeps horizontally (along the x-axis) over a set of $n$ entry areas. To store which areas overlap on the y-axis, we use two $n$ sized vectors, called counting vector $V_c$ and pointer vector $V_p$. $V_c$ and $V_p$ are initialized with zeros and null pointers, respectively. During the line-sweep, $V_c$'s purpose is to hold the number of overlapping entry areas for each y-position, while the pointers in $V_p$ shall point to the clusters of entry areas that occupy the respective y-positions. To map the y-values of all entry areas to the indices of the two vectors, we apply y-squeezing to all entry areas. y-squeezing is achieved by creating a sorted list of all y-values (top and bottom of all entry-areas) and using each y-value's index in the sorted list as the index for $V_c$ and $V_p$. The line sweep iterates over the x-values of the start and end-points of all entry areas in ascending order. At the start point of each area, we merge this area with all overlapping clusters (stored in $V_p$). At the end-point of each area, we check (using $V_c$) if we can remove the area's cluster from $V_p$. In detail, this is performed as follows:

- Whenever the line sweep stops at the start position of an entry area $e = (e_{top}, e_{bottom})$, we increase all counters in $V_c[e_{top}, e_{bottom}]$. Further, let $C$ be the set of clusters pointed to in $V_p[e_{top}, e_{bottom}]$. We create a new set $S$, that contains all entries in $C$ as well as $e$. Let $P$ be the set of pointers in $V_p$ that point to any cluster in $C$. (Note that $P$ contains all pointers in $V_p[e_{top}, e_{bottom}]$, but may contain further pointers.) All pointers in $P$ are redirected to $S$. This redirection does not require a scan over the full vector, since we memorize the maximal y-axis interval of each cluster. However, before redirecting a pointer in the y-axis interval of a cluster, we need to verify that the pointer is not pointing to another cluster, as a cluster can be hollow.
- Whenever the line-sweep stops at the end position of an entry area $e = (e_{top}, e_{bottom})$, we decrease all entries in $V_c[e_{top}, e_{bottom}]$. For each value in $V_c$ that becomes zero, we set the respective pointer in $V_p$ to null. We detect if this step removes the last pointer to a cluster by memorizing the number of active pointers for each cluster. Any cluster that becomes pointer-less is saved as the output of the line sweep.

#### The inexact (heuristic) algorithm

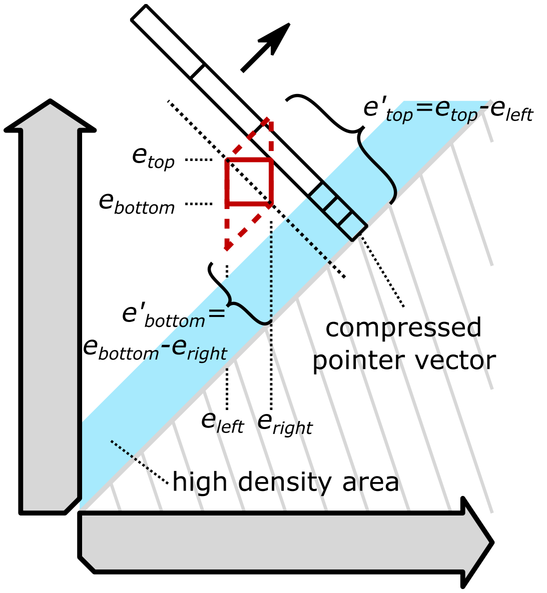

The exact algorithm requires memory relative to the size of $V_c$ and $V_p$, which is too large for big genomes. Hence, the inexact line sweep uses a compressed form of the pointer vector and drops the counting vector. This compression is achieved by having each counter represent an interval instead of a single point on the y-axis. However, there is a high density of entries just above the diagonal, where a compressed vector would lead to the creation of a single genome-spanning cluster. To avoid this genome-spanning cluster, we do not compress $V_p$ in this area of high density. Since the area of high-density moves along the diagonal (i.e. upwards on the y-axis) during the line sweep, we perform a coordinate system transformation on the y-coordinates of all entries. With the transformed y-coordinates, the high-density area remains in the same interval for the entire line-sweep. We compute the new y-coordinates $e'_{top}$ and $e'_{bottom}$ for each entry area $e$ as follows:

$$e'_{top} = e_{top} - e_{left}$$

$$e'_{bottom} = e_{bottom} - e_{right}$$

Note that this transformation turns all rectangles into parallelograms in the original coordinate system. By placing all parallelograms so that they surround their original rectangles, we ensure that exact clusters of rectangles end up in the same inexact cluster.

Instead of using $V_c$ to track the exact extension of each set on the y-axis, we use the memorized maximal y-axis interval as well as a counter of open entry areas for each set. This saves us from adjusting $V_c$ but leads to clusters casting "shadows" to their right. (This shadow ends at the endpoint of the rightmost parallelograms in the cluster.) Clusters that are erroneously joined due to the “shadows” or the coordinate transformation are separated again during the exact line-sweep.

#### Supplementary Note 14. Reconstructing sequenced genomes from graphs

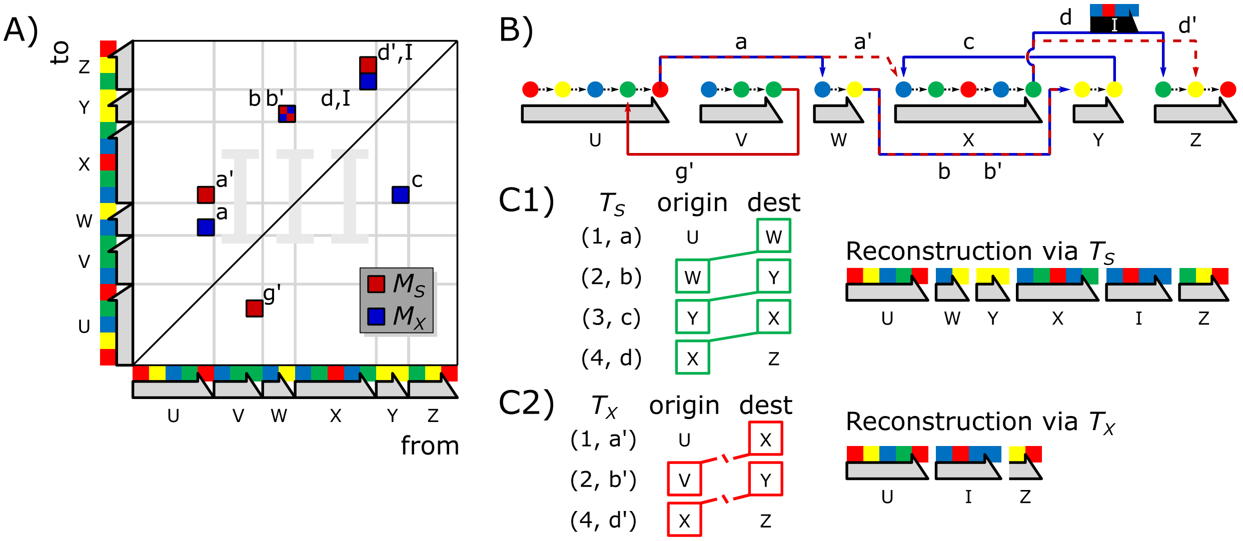

The figure visualizes an example of the reconstruction of a sequenced genome $S = UWYXIZ$ from a reference genome $UVWXYZ$. The reconstruction happens via a single error-free read spanning $S$ (subfigure C1) and via a set $X$ of simulated long reads (PacBio) and short reads (Illumina) for $S$ (subfigure C2). **A)** shows the third quadrant of the two adjacency matrices $M_S$ and $M_X$ for C1) and C2) in combined form, respectively. Here the red entries belong to $M_S$ and the blue entries belong to $M_X$. $(a, a')$, $(b, b')$ and $(d, d')$ represent pairs of spatially closest neighbors in $M_X$ and $M_S$. The entry $c \in M_S$ does not have a corresponding entry in $M_X$. Similarly, the entry $g' \in M_X$ is without a partner in $M_S$. Subfigure **B)** displays the graphs $G_S$ and $G_X$ for $M_S$ and $M_X$ in combined form. (Red edges are part of $G_S$ and belong to $M_S$, while blue edges are part of $G_X$ and belong to $M_X$.) Here the weight $I$ of the edges $d$ and $d'$ represents a sequence not occurring on the reference genome that is inserted whenever the reconstruction passes through $d$ or $d'$. The tables in **C1)** and **C2)** list the elements of the sets $T_S$ and $T_X$, respectively. $T_S$ and $T_X$ describe traversals through their respective graphs, as explained in the main text. Additionally, the tables in C1) and C2) annotate the origin and destination of each row's edge. The genomes that are reconstructed from the tables are shown to their right. The tables additionally annotate matching (green lines) and non-matching (red lines) connections between the 'origin' and 'destination' entries of consecutive rows.

#### Supplementary References

1. Schmidt M, Heese K, Kutzner A: **Accurate high throughput alignment via line sweep-based seed processing**. *Nature Communications* 2019, **10**(1):1939.
2. Abadi M, Barham P, Chen J, Chen Z, Davis A, Dean J, Devin M, Ghemawat S, Irving G, Isard M: **Tensorflow: A system for large-scale machine learning**. In: *12th {USENIX} symposium on operating systems design and implementation ({OSDI} 16): 2016*. 265-283.
3. Douglas K, Douglas S: **PostgreSQL**: New Riders Publishing; 2003.
4. Groff J, Weinberg P: **SQL The Complete Reference, 3rd Edition**: McGraw-Hill, Inc.; 2009.
5. **Bokeh: Python library for interactive visualization**. 2020.
